# WNT inhibition creates a BRCA-like state in Wnt-addicted cancer

**DOI:** 10.1101/2020.06.17.157024

**Authors:** Amanpreet Kaur, Sugunavathi Sepramaniam, Jun Yi Stanley Lim, Siddhi Patnaik, Nathan Harmston, May Ann Lee, Enrico Petretto, David M. Virshup, Babita Madan

## Abstract

Wnt signaling maintains diverse adult stem cell compartments and is implicated in chemotherapy resistance in cancer. PORCN inhibitors that block Wnt secretion have proven effective in Wnt-addicted preclinical cancer models and are in clinical trials. In a survey for potential combination therapies, we found that Wnt inhibition synergizes with the PARP inhibitor olaparib in Wnt-addicted cancers. Mechanistically, we find that multiple genes in the homologous recombination and Fanconi anemia repair pathways, including *BRCA1*, *FANCD2*, and *RAD51* are dependent on Wnt/β-catenin signaling in Wnt-high cancers, and treatment with a PORCN inhibitor creates a BRCA-like state. This coherent regulation of DNA repair genes occurs via a Wnt/β-catenin/MYBL2 axis. Importantly, this pathway also functions in intestinal crypts, where high expression of BRCA and Fanconi anemia genes is seen in intestinal stem cells, with further upregulation in Wnt high APC^min^ mutant polyps. Our findings suggest a general paradigm that Wnt/β-catenin signaling enhances DNA repair in stem cells and cancers to maintain genomic integrity. Conversely, interventions that block Wnt signaling may sensitize cancers to radiation and other DNA damaging agents.

## INTRODUCTION

Stem cells in normal tissues protect the integrity of their genome by expression of diverse proteins that accurately detect and repair mutations introduced by DNA replication and environmental mutagens (1). Inherited defects in DNA repair pathways, such as germline mutations in BRCA1 or Fanconi anemia pathway genes that normally repair double-strand breaks and DNA crosslinks, lead to the accumulation of mutations that impair stem cell function and promote tumorigenesis (2, 3). The FA pathway proteins serve as interstrand crosslink (ICL)-sensors and promote DNA repair in conjunction with homologous recombination and other DNA repair pathways including BRCA proteins (4, 5). Individuals with Fanconi anemia who have mutations in Fanconi anemia (FA) pathway genes are exquisitely sensitive to DNA ICL-generating agents. These DNA repair pathways are often co-opted in cancers during the development of drug resistance (6).

Defects in specific DNA repair pathways also present an opportunity for targeted anti-cancer therapies (2, 3, 7). Breast and ovarian cancers with defects in the homologous recombination (HR) repair pathway, especially BRCA1 and BRCA2 mutant tumors, are sensitive to poly ADP-ribose polymerase (PARP) inhibitors such as olaparib (7–9). These inhibitors work by blocking the function of PARP proteins essential for single-strand break (SSB) repair. Upon PARP inhibitor treatment, the unrepaired SSBs can progress to double-strand breaks that are then repaired by homologous recombination. In HR-deficient cells, treatment with PARP inhibitors results in growth arrest by apoptosis or senescence due to the accumulation of DNA damage (7).

Wnt signaling is important for the maintenance of stem cell state, adult tissue homeostasis and the prevention of differentiation (10). Wnt signaling is also associated with the development of radioresistance in many cancers but the underlying mechanisms are not well understood (11–16). Wnts are a family of 19 secreted palmitoleated glycoproteins that signal by binding to Frizzled (FZD) and additional co-receptors on the cell surface. The interaction of Wnts with their receptors stabilizes β-catenin, which then translocates to the nucleus to regulate gene expression. Aberrant stabilization of β-catenin can be caused by mutation of components of the Wnt pathway such as *APC*, *AXIN2* and *CTNNB1* (β-catenin), as is frequently seen in colorectal, gastric and liver cancers (17). Additionally, a subset of pancreatic, mucinous ovarian, colorectal, gastric, adrenocortical and endometrial cancers harbor loss-of-function mutations in E3-ubiquitin ligases *RNF43* or its paralog *ZNRF3*, or gene fusions leading to activation of Wnt agonists *RSPO2/3* (R-spondin 2/3) (18–24). These mutations enhance the abundance of Frizzleds and cause cancers to be dependent on activated Wnt signaling. This subset of cancers is highly sensitive to upstream inhibitors of Wnt signaling pathway such as anti-FZD and anti-R-spondin antibodies as well as PORCN inhibitors such as ETC-159 (25–27). Several of these agents are currently in clinical trials. These inhibitors are also powerful tools to investigate the pathways that are regulated by Wnt signaling in cancer (27, 28).

During a screen to identify approved drugs that synergize with PORCN inhibitors, we made the unexpected observation that the PARP inhibitor olaparib synergized with ETC-159 in Wnt-addicted cancers. Mechanistically, we found that inhibition of Wnt signaling in multiple cancer cell lines and normal intestinal crypts results in the suppression of multiple HR pathway genes. Wnt signaling through a β-catenin/MYBL2 pathway regulates the expression of *BRCA1*, *BRCA2*, *RAD51* and *FANCD2*. This study uncovers a role for Wnt/β-catenin signaling in the regulation of homologous recombination DNA repair pathway in intestinal stem cells and in cancer, and demonstrates that inhibition of Wnt signaling confers a BRCA-like phenotype, providing a novel therapeutic opportunity for Wnt high cancers.

## RESULTS

### Wnt inhibition synergizes with PARP inhibitor Olaparib

The PORCN inhibitor ETC-159 has shown efficacy as a monotherapy in preclinical models of Wnt-addicted cancers (27). To identify potential combinatorial therapeutic options, we performed a synergy screen using the Chou-Talalay method with selected drugs that are either FDA approved or in clinical trials (29). Since Wnt-addicted cells are substantially more sensitive to PORCN inhibitors when grown in suspension, the screen was performed in the Wnt-addicted RNF43-mutant pancreatic cancer cell line HPAF-II using soft agar colony formation as the readout (27, 30). We observed that the combination of olaparib and ETC-159 was significantly more effective in inhibiting colony formation of HPAF-II cells than treatment with either drug individually (Figure 1A and Table S1). Consistent with this, the drug combination index values for ETC-159 and olaparib, as determined using the Chou-Talalay CompuSyn algorithm, showed a synergistic effect (Combination Index < 1) for each of the dosages tested (Figure 1B). To test if this combination was also efficacious *in vivo*, we used the HPAF-II pancreatic cancer xenograft model. Treatment with olaparib alone or low dose of ETC-159 alone was less effective compared to the combination of olaparib and ETC-159 in preventing HPAF-II tumor growth in mice. This was shown by changes in tumor volumes during the course of treatment (Figure 1C) as well as the tumor weights at the end of 21 days treatment (Figure 1D).

**Figure 1:**
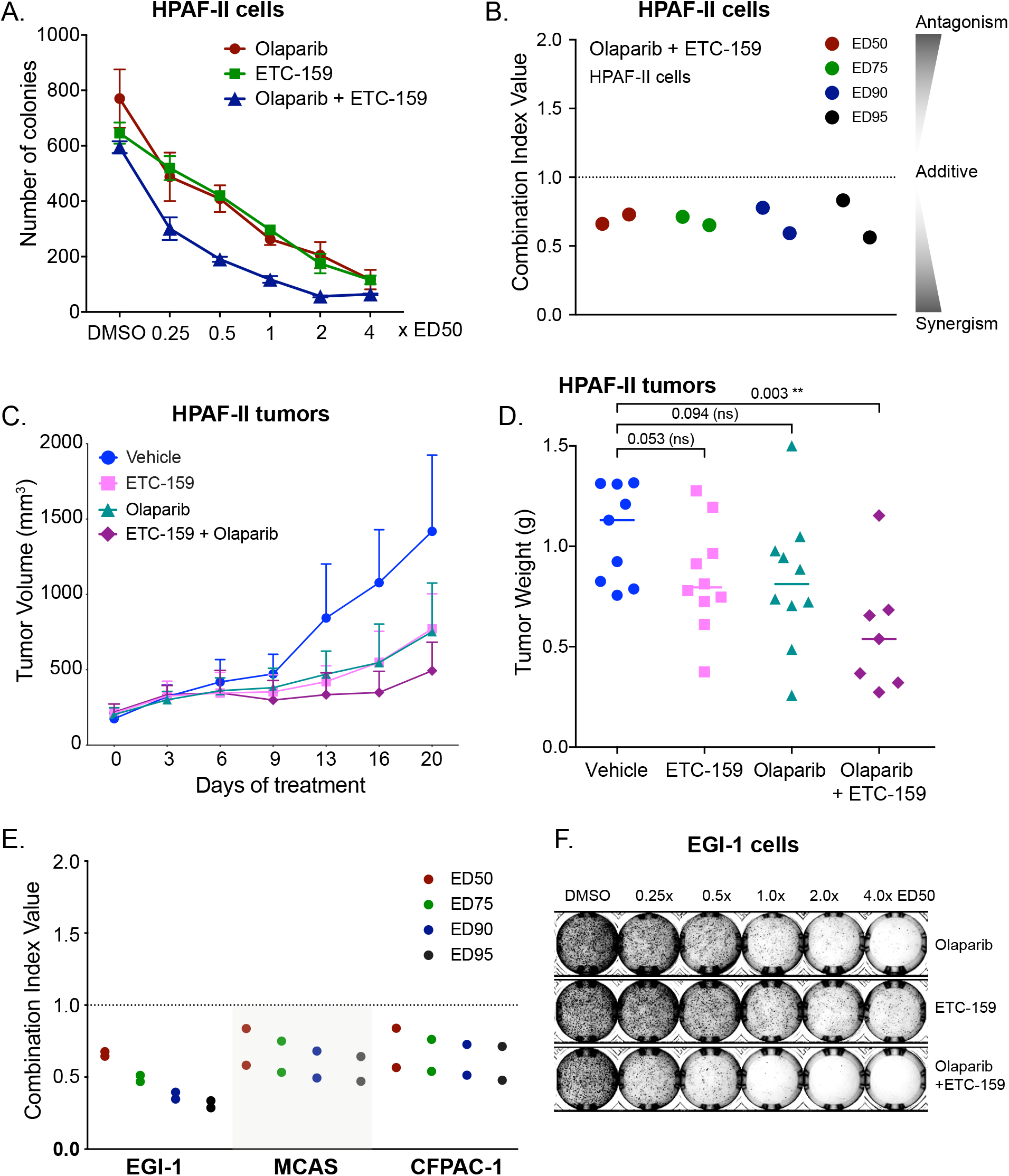
ETC-159 synergizes with the PARP inhibitor Olaparib. **A-B**. ***PORCN inhibitor ETC-159 synergizes with PARP inhibitor olaparib in suppressing proliferation of HPAF-II pancreatic cancer cells in soft agar.*** The ED_50_ (dose that reduced colony formation by 50% of the maximal inhibition) was determined (Table S1) and cells were treated with ETC-159, olaparib or the combination at the indicated dose (for example 0.25 × ED_50_ of Olaparib or ETC-159, or 0.25 × ED_50_ of Olaparib + 0.25 × ED_50_ of ETC-159, respectively). (A) The data is representative of two independent experiments with each point representing an average colony count ± SD of duplicates. (B) The Combination Index (CI) values of olaparib and ETC-159 calculated for the two independent experiments using the Chou-Talalay CompuSyn software. CI < 1, = 1, and > 1 indicate synergism, additive effect, and antagonism, respectively. The lower the CI, the stronger the synergism. **C-D**. ***Olaparib and ETC-159 synergize to prevent the growth of HPAF-II xenografts in mice.*** NSG mice with established HPAF-II subcutaneous xenografts were randomized into four groups. Mice were gavaged daily with ETC-159 (10 mg/kg), Olaparib (50 mg/kg) or a combination of ETC-159 (10 mg/kg) and Olaparib (50 mg/kg). Treatment was initiated after HPAF-II tumors were established. (C) Tumor volumes were measured starting from day 0 and during the course of treatment as shown. Data points represent the mean ± SD. n = 7-8 tumors/group. (D) Tumor weights in the respective groups at the end of the treatment are shown, each dot represents an individual tumor. p-values were calculated with Mann-Whitney test. **E. *Olaparib and ETC-159 synergize in multiple Wnt-addicted cancer cells.*** Soft agar colony formation assays were performed as in Figure 1A with the indicated cell lines treated with varying concentrations of ETC-159, olaparib or a combination of both as indicated and the combination index was calculated. PARP inhibitor olaparib and ETC-159 synergistically prevent colony formation of EGI-1, MCAS and CFPAC-1 cells in soft agar. **F.** Representative image of soft agar colonies of EGI-1 cells is shown.

Next, we assessed the combination of PORCN inhibitor and olaparib in three additional Wnt-addicted cell lines from diverse cancer types with distinct Wnt pathway mutations. The cholangiocarcinoma cell line EGI-1 (Wnt-addicted due to an R-spondin translocation), the ovarian cancer cell line MCAS (Wnt-addicted due to an *RNF43* mutation), and the pancreatic cancer cell line CFPAC-1 (sensitive to Wnt inhibition, mechanism unknown) were used. Similar to what was observed with HPAF-II cells, the combination of ETC-159 and olaparib synergistically inhibited colony formation in all three cell lines in soft agar assay at all the doses tested (Figures 1E-F and Table S1). Thus, the synergy of ETC-159 and olaparib is a general phenomenon. Taken together, the data indicates that blocking Wnt activity with a PORCN inhibitor sensitizes Wnt-addicted cells to a PARP inhibitor.

### Wnt inhibition reduces expression of homologous recombination (HR) and Fanconi anemia (FA) repair pathway genes

Olaparib and related PARP inhibitors are uniquely effective in BRCA-mutant and BRCA-like cancers that have dysfunctional homologous recombination (31). Diverse mechanisms can cause defective BRCA-like behavior, including inherited mutations in *BRCA1*, *BRCA2* and Fanconi anemia complementation (*FANC*) group of genes (32). Epigenetic and transcriptional mechanisms that silence HR pathway genes can also cause a BRCA-like state (33). To test if Wnt inhibition induces a BRCA-like state, we examined the expression of genes involved in DNA repair using our transcriptome dataset from ETC-159 treated Wnt-addicted HPAF-II pancreatic cancer orthotopic xenografts (Figure S1A) (28). Remarkably, amongst the genes downregulated following PORCN inhibition, i.e *Wnt-activated* genes, there were three clusters of genes (C1, C5 and C12) that were significantly enriched for Gene Ontology (GO) annotated processes and pathways related to multiple components of the DNA damage repair pathway (Figures 2A-B and S1A).

**Figure 2:**
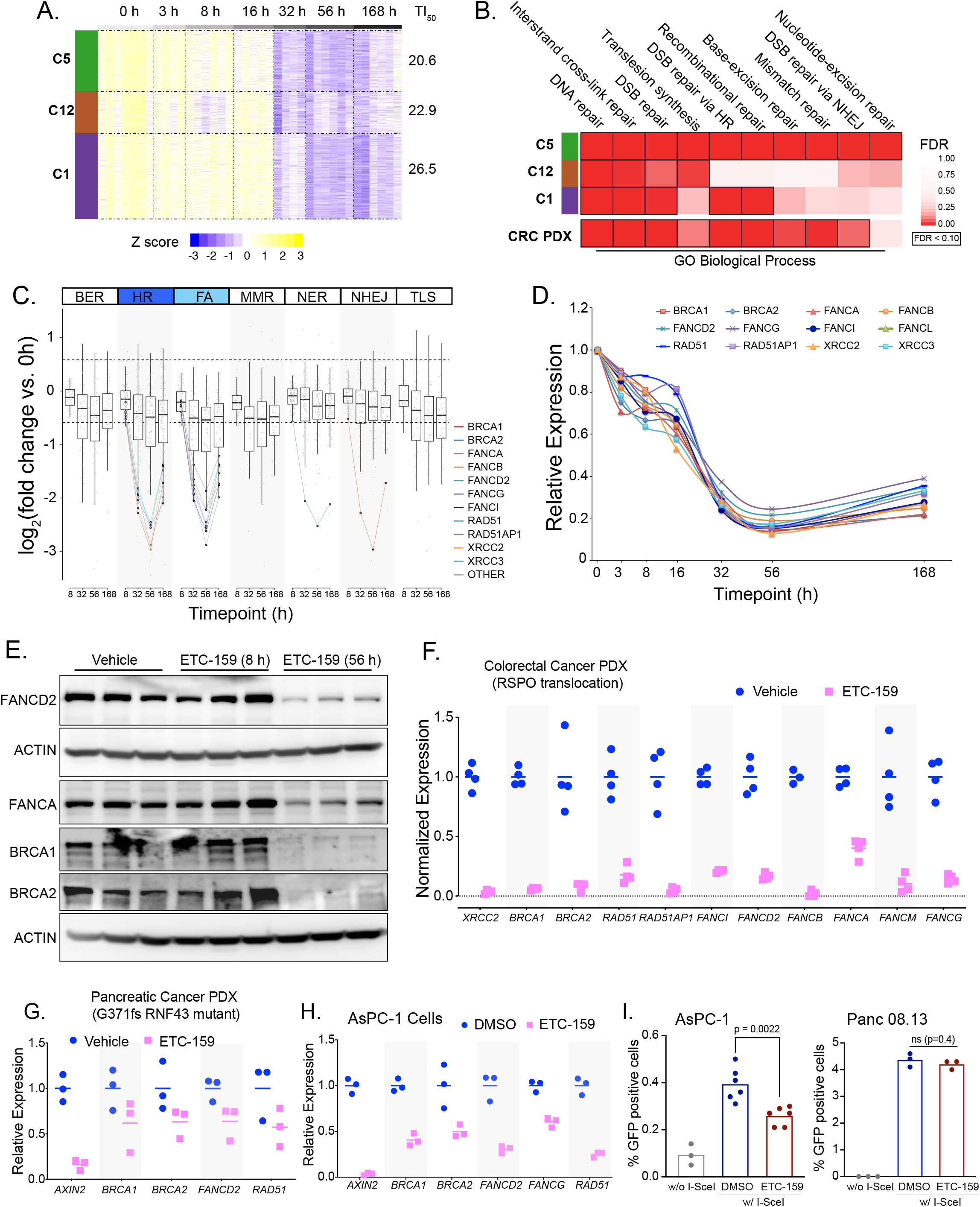
Homologous recombination (HR) and Fanconi anemia (FA) repair pathway genes are regulated by Wnt signaling. **A. *Heatmap of selected temporal clusters containing the Wnt-activated genes that are enriched for DNA repair pathways.*** Transcriptomic data from HPAF-II orthotopic pancreatic tumors (dataset originally reported in 28)) was assessed at multiple time points during treatment with the PORCN inhibitor 37.5 mg/kg *bid* ETC-159 (Figure S1A). Genes that were differentially expressed over time (FDR < 10%) following PORCN inhibition were clustered based on their pattern of transcriptional response. Clusters 1, 5 and 12 refer to temporal clusters defined in (28) and TI_50_ is the time (in hours) after start of therapy to achieve 50% inhibition in gene expression for each cluster. **B. *Gene Ontology (GO) biological process enrichment of each cluster of Wnt-activated genes.*** Analysis of *Wnt-activated* genes in clusters 1, 5 and 12 from HPAF-II orthotopic pancreatic tumors (Figure 2A) and colorectal cancer (CRC) patient-derived xenograft (PDX) highlights enrichment of genes involved in multiple DNA repair pathways such including interstrand cross-link repair, double strand break (DSB) repair via homologous recombination (HR). **C. *DNA repair genes involved in HR and FA pathways were differentially expressed over time in response to PORCN inhibition in HPAF-II orthotopic xenografts.*** Comparison of log2 fold change (FDR < 10%) in the expression of *Wnt-activated* genes over time across multiple DNA repair pathways shows that genes regulating HR and FA pathways have higher fold changes compared to genes regulating BER, NER or other pathways. BER: base excision repair; HR: homologous recombination; FA: Fanconi anemia; MMR: mismatch repair; NER: nucleotide excision repair; NHEJ: non-homologous end joining; TLS: translesion DNA synthesis. **D. *ETC-159 treatment of HPAF-II orthotopic xenografts reduces the expression of multiple HR and FA pathway genes.*** Temporal expression of selected *Wnt-activated* HR and FA pathway genes are shown. n = 7-10 tumors/group. **E. *Protein levels of HR and FA pathway genes are reduced upon Wnt inhibition.*** Tumor lysates from HPAF-II xenografts treated with vehicle or ETC-159 for 8 or 56 hours were analyzed by SDS-PAGE and immunoblotted with the indicated antibodies. Each lane represents an individual tumor. **F. *Wnt inhibition reduces the expression of HR and FA pathway genes in a Wnt-addicted colorectal cancer patient-derived xenograft.*** Mice with CRC PDX driven by a *RSPO3* translocation were treated with vehicle or ETC-159 for 56 hours before harvesting and analyzed by RNA-seq (dataset reported in (27)). The graph shows the normalized expression of the indicated genes. n = 4 tumors/group. **G. *Wnt inhibition reduces the expression of HR and FA pathway genes in a RNF43-mutant pancreatic cancer patient-derived xenograft.*** Pancreatic cancer PDX with G371fs RNF43 mutation treated with vehicle or ETC-159 (30 mg/kg) for 21 days were analysed for changes in the expression of the indicated genes measured by qRT-PCR. Each data point represents an individual tumor. n = 3 tumors/group. **H. *Wnt inhibition reduces the expression of HR and FA pathway genes in RNF43-mutant AsPC-1 cells.*** AsPC-1 cells were seeded in low adherence plates, treated with DMSO or ETC-159 (100 nM) for 72 hours and then the expression of the indicated genes was measured by qRT-PCR. **I. *Wnt inhibition reduces homologous recombination***. AsPC-1 or Panc 08.13 cells with a stable integration of HR reporter construct DR-GFP (encoding a modified GFP gene with the I-SceI site and a donor GFP sequence that restores GFP sequence upon HR-mediated DNA repair) were transfected with a plasmid expressing I-SceI endonuclease, followed by treatment with DMSO or ETC-159 (100 nM) for 24 hours. The percentage of GFP positive cells as measured by flow cytometry indicates that the extent of repair via HR pathway in the presence of ETC-159 treatment is reduced in Wnt high AsPC-1 cells or but not in Wnt-insensitive Panc 08.13 cells. Data shows the values from three independent experiments with two to three replicates each.

Further analysis of the genes involved in regulating each of the DNA repair pathways highlighted that Wnt inhibition robustly downregulated genes involved in homologous recombination (HR) and Fanconi anemia (FA) pathways (Figure 2C). By comparison, genes annotated to be involved in mismatch repair and in the repair of single strand breaks such as base excision repair (BER) or nucleotide excision repair (NER) were much less affected by changes in Wnt signaling. Most notably, the expression of *BRCA1*, *BRCA2*, *RAD51* and multiple *FANC* genes was decreased ∼8-10 fold by 32 hours after the start of treatment with the PORCN inhibitor (Figures 2C-D). Immunoblotting for BRCA1, BRCA2, FANCA and FANCD2 in HPAF-II tumors harvested 8 and 56 hours after the start of ETC-159 treatment confirmed the downregulation of these proteins (Figure 2E). This downregulation of genes involved in the HR and FA pathways following Wnt inhibition is consistent with the observed increase in sensitivity of Wnt-addicted tumors and cell lines to olaparib (Figure 1).

While ETC-159 treatment of HPAF-II cells *in vitro* for 48 h significantly reduced the expression of HR-pathway and FA genes, we did not observe a significant change in the percentage of cells in S-phase as compared to control (Figures S1B-C). These results suggest that the suppression of HR and FA pathway gene expression by ETC-159 treatment is not due to its effect on the cell proliferation (34, 35).

To test if PORCN inhibition causes a decrease in HR and FA genes in additional Wnt-addicted cancer models, we examined RNA-seq data from a colorectal cancer patient-derived xenograft (CRC PDX) with a *RSPO3* translocation that was treated with ETC-159 (27). In this CRC PDX model, similar to the HPAF-II xenografts, genes that were downregulated (FDR < 0.1, fold-change > 1.5) also had significant enrichment for multiple processes associated with DNA damage repair (Figure 2B). Specifically, several key HR and FA pathway genes were downregulated upon Wnt inhibition (Figure 2F) (27, 28). Consistent with this being a general phenomenon in Wnt-addicted cancers, HR pathway gene expression as analyzed by qRT-PCR was also decreased upon ETC-159 treatment in a pancreatic cancer patient-derived xenograft with an *RNF43* mutation (G371fs), as well as in AsPC-1 pancreatic cancer and EGI-1 cholangiocarcinoma cells (Figures 2G-H and S1D).

A global decrease in multiple DNA repair enzymes including BRCA1 and RAD51 prevented our ability to measure strand breakage using standard assays such as foci formation. Therefore, to test the functional impact of reduced HR pathway gene expression on homologous recombination we used an established assay for homology-directed repair, the Direct Repeats (DR)-GFP assay (36). We compared the effect of Wnt inhibition on GFP recombination using AsPC-1 cells. AsPC-1 cells with stably integrated DR-GFP reporter construct containing a modified GFP gene disrupted by an I-SceI cleavage site and a donor GFP sequence that restores functional GFP upon HR-mediated DNA repair. In DR-GFP AsPC-1 cells, expression of the I-SceI endonuclease led to a four-fold increase in the percentage of GFP positive cells. However, Wnt inhibition with ETC-159 for 48 hours led to ∼40% reduction in the percentage of cells undergoing homology-directed repair (GFP positive) (Figure 2I). As a control for off-target effects of ETC-159, we performed the same assay in Panc 08.13 cells that do not have pathologic activation of Wnt signaling. In these cells, ETC-159 treatment had no effect on the recombination repair efficiency of as measured by percentage of GFP positive cells. (Figure 2I).

Taken together, the data from multiple models of Wnt-addicted cancers demonstrates that inhibition of Wnt signaling sensitizes cells to the PARP inhibitor olaparib by creating a BRCA-like state via downregulation of HR and FA pathways.

### Wnt high cells that are resistant to PORCN inhibition can be sensitized by combination with olaparib

There is a subset of cells with high Wnt signaling whose growth is not slowed down by PORCN inhibition alone (26). PaTu8988T cells are Wnt high owing to a loss-of-function RNF43 mutation (F69C), have high abundance of Frizzleds on the cell surface, and show reduced expression of Wnt target genes upon knockdown of β-catenin showing that the regulation of these genes is Wnt regulated (Figures 3A-B). However, their proliferation is relatively insensitive to Wnt inhibition alone, presumably due to a second downstream mutation (26). Despite their continued proliferation, PaTu8988T cells showed downregulation of both HR and FA pathway genes in response to Wnt inhibition with either siβ-catenin or ETC-159 treatment (Figure 3B). A high dose of ∼1.0 μM of either ETC-159 or olaparib reduced colony formation by only 50% (Figure 3C and Table S1), with no further reduction even when the dose was increased up to 4.0 μM (4.0× ED_50_). However, the combination of olaparib and ETC-159 was highly synergistic even at 250 nM (0.25× ED_50_) (Combination Index < 0.2 at three dosages tested) (Figures 3C-D). This is consistent with ETC-159 creating a BRCA-like state via inhibition of the Wnt pathway rather than through an off-target effect. Importantly, these data suggest that the combination of olaparib and ETC-159 may be an effective treatment for some Wnt high cancers with inactivating RNF43 mutations that are resistant to monotherapy with Wnt pathway inhibitors.

**Figure 3:**
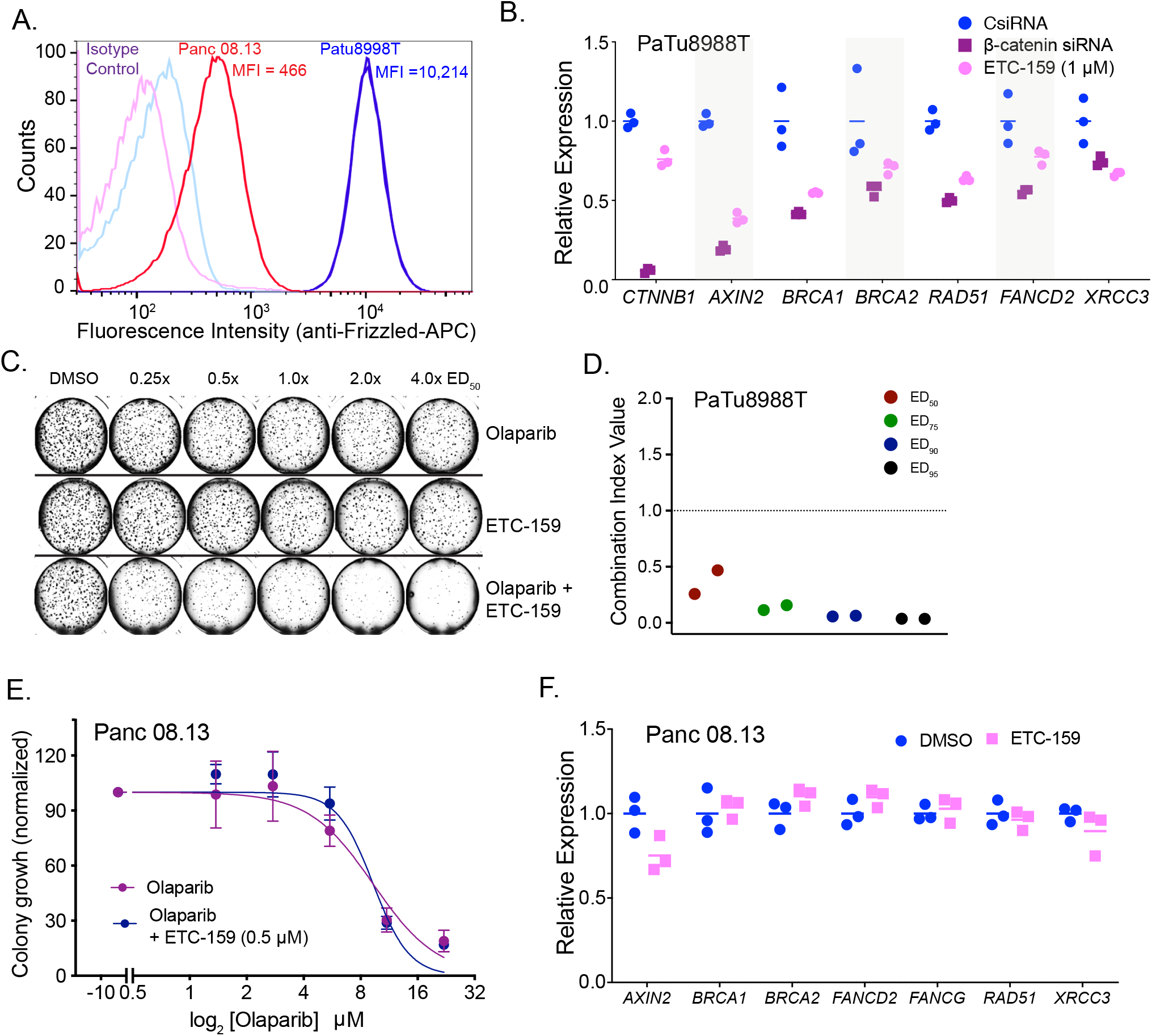
Wnt high cells that are resistant to PORCN inhibition can be sensitized by combination with olaparib. **A. *Flow cytometric analysis of endogenous cell surface FZD levels in PaTu8988T cells with inactivating RNF43 mutation (F69C) shows high cell surface abundance of FZDs.*** PaTu8988T cells were stained with pan-FZD antibody clone F2.A followed by anti-human Fc fragment APC-conjugated secondary antibody. MFI = median fluorescence intensity. **B. *Wnt inhibition using siRNA against β-catenin or ETC-159 treatment reduces the expression of HR and FA pathway genes in PaTu8988T cells.*** PaTu8988T cells were transfected with indicated siRNAs or treated with ETC-159 (1 *μ*M) for 48 hours, total RNA was isolated and expression of β-catenin (*CTNNB1*), *AXIN2* and DNA repair genes was measured by qRT-PCR. **C-D. *ETC-159 and olaparib synergistically inhibit the growth of ETC-159 resistant pancreatic cancer cell line PaTu8988T.*** (C) PaTu8988T cells were seeded in soft agar and treated with ETC-159, olaparib or the combination of the two inhibitors at an equivalent dose as described in Figure 1A. Representative image of soft agar colonies from PaTu8988T cells combination study in a 48-well plate is shown. (D) The Combination Index (CI) values of olaparib and ETC-159 calculated from two independent experiments using the Chou-Talalay CompuSyn software. **E. *Flow cytometric analysis of endogenous cell surface FZD levels in Panc 08.13 cells with wild-type RNF43 shows low cell surface abundance of FZDs.*** Panc 08.13 cells were stained as described in Figure 3A followed by flow cytometric analysis. MFI = median fluorescence intensity. **F. *Wnt inhibition does not synergize with olaparib in Wnt-low Panc 08.13 cells*.** Panc 08.13 cells were seeded in soft agar and treated with ETC-159, olaparib or the combination of the two inhibitors at the indicated doses. The data is representative of two independent experiments with each point representing an average colony count ± SD of duplicates. **G. *Wnt inhibition through ETC-159 does not change the expression of HR and FA pathway genes in Panc 08.13 cells.*** Panc 08.13 cells were cultured in low adherence plates and treated with ETC-159 for 72 hours, total RNA was isolated and expression of indicated genes was measured by qRT-PCR.

On the other hand, Panc 08.13, a pancreatic cancer cell line with no RNF43 mutation has low abundance of Frizzleds on the cell surface and hence has low Wnt signaling (Figure 3A). Consistent with low Wnt signaling, Panc 08.13 cells were not sensitive to Wnt inhibition and no synergy was observed between ETC-159 and olaparib in soft agar colony formation assay (Figure 3E). Panc 08.13 cells also did not show a reduction in the expression of HR and FA pathway genes upon ETC-159 treatment (Figure 3F).

### Regulation of DNA repair genes is β-catenin dependent in Wnt-addicted cells

Wnt signaling regulates gene expression in large part through stabilization of β-catenin (37). To test if HR and FA pathway gene expression was β-catenin dependent, we knocked down β-catenin expression in HPAF-II cells with siRNA. Similar to Wnt inhibition with ETC-159, knockdown of β-catenin in HPAF-II cells caused a marked decrease in the expression of multiple HR and FA pathway genes (Figure 4A). Conversely, we tested if degradation of β-catenin was required for the downregulation of these genes. We established xenograft tumors using either HPAF-II cells, or HPAF-II cells that express a stabilized β-catenin that is insensitive to CK1α/GSK3 phosphorylation and subsequent proteasomal degradation. In these latter tumors, treatment with ETC-159 should not impact the expression of Wnt/β-catenin target genes. This also serves as a control for determining if the downstream effects are due to off-target effects of the PORCN inhibitor. Tumor-bearing mice were treated with ETC-159 for 56 hours, and then the expression of HR and FA pathway genes were compared between the two groups. As expected, ETC-159 reduced the expression of HR pathway genes in the control HPAF-II tumors, but this effect was totally abrogated in the stabilized β-catenin group tumors (Figure 4B). Moreover, in a soft agar colony formation assay, we observed that unlike the HPAF-II cells (Figure 1A), co-treatment with ETC-159 failed to sensitize HPAF-II cells with stabilized β-catenin to olaparib (Figure 4C). These findings confirm the importance of β-catenin signaling in the regulation of DNA repair genes.

**Figure 4:**
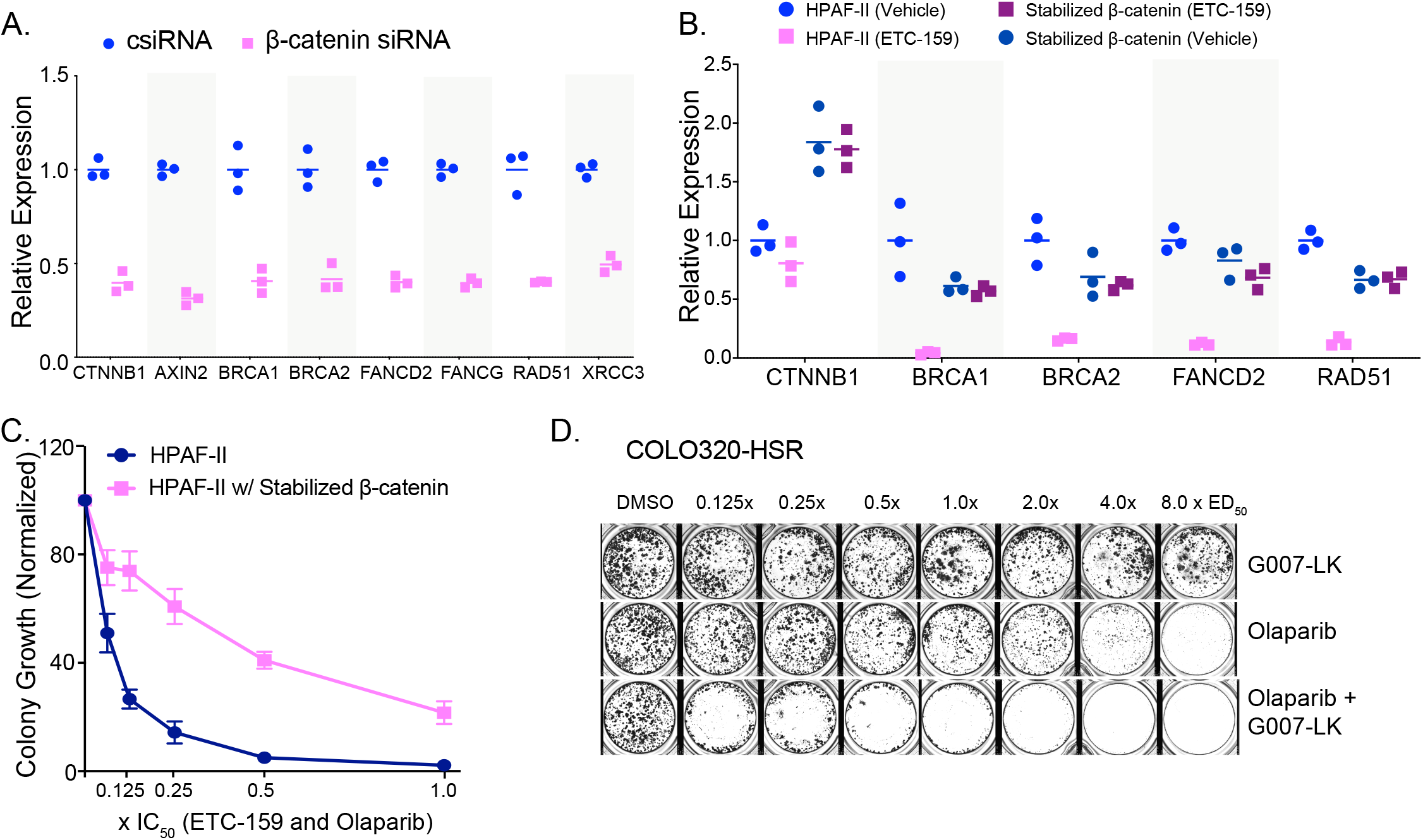
β-catenin regulates the expression of HR and FA pathway genes. **A. *β-catenin regulates the expression of HR and FA pathway genes in HPAF-II cells***. HPAF-II cells were transfected with siRNA against β-catenin for 48 hours. The relative expression of genes as measured by qRT-PCR is shown. **B. *Stabilized β-catenin prevents the downregulation of HR and FA pathway genes upon Wnt inhibition.*** Mice bearing HPAF-II xenografts without or with stabilized β-catenin were treated with ETC-159 (37.5mg/kg *bid*) for 56 hours before tumors were harvested and the expression of indicated genes was measured by qRT-PCR. The ETC-159 induced reduction in the expression of HR and FA pathway genes was blocked in xenografts with stabilized β-catenin. **C. *Stabilized β-catenin reduces the sensitivity of HPAF-II cells to olaparib and ETC-159.*** HPAF-II cells without or with stabilized β-catenin were plated in soft agar in 48 well plates. The cells were treated with a combination of olaparib and ETC-159 as in Figure 1A and the total number of colonies were scored after 2 weeks. **D. *G007-LK and olaparib synergistically inhibit the growth of COLO320-HSR colorectal cancer cells with APC mutation***. COLO320-HSR cells were plated at a low density and treated with G007-LK, olaparib or the combination of the two inhibitors at an equivalent dose as described in Figure 1A. Representative image of two independent experiments is shown.

Finally, we tested if a drug that inhibits Wnt/β-catenin signaling via an entirely different mechanism also regulated HR and FA pathway. We selected G007-LK, a small molecule that inhibits Wnt signaling by stabilizing the destruction complex protein Axin, hence facilitating the degradation of β-catenin (38). G007-LK is selective for TNKS1/2 (IC_50_ < 50nM) and does not inhibit PARP1/2 (IC_50_ > 10μM) at therapeutic doses (38). The APC mutant colorectal cancer cell line, COLO320-HSR was treated with increasing concentrations of G007-LK alone or in combination with olaparib (Figure 4D). While G007-LK and olaparib alone were not particularly effective in preventing the growth of these cells, the combination of the two drugs was synergistic in inhibiting the growth of COLO320 HSR cells (Combination Index of 0.3 at ED_50_, 0.17 at ED_75_, 0.098 at ED_90_ and 0.072 at ED_95_). These results demonstrate that treatment with two independent Wnt inhibitors sensitizes Wnt-high cells to olaparib.

### Identification of a Wnt/β-catenin/MYBL2 pathway

To understand the mechanism of regulation of HR pathway genes by Wnt/β-catenin signaling, we analyzed the promoters of the HR and FA pathway genes that were coherently downregulated upon Wnt inhibition. Transcription factor binding site (TFBS) targets were obtained from the ENCODE, ReMAP, and Literature ChIP-seq datasets using the online tool CHEA3 (39). This analysis identified significant enrichment for FOXM1 and MYBL2 binding events (Figure 5A) which is consistent with the established roles of these two transcription factors in the regulation of DNA repair pathways (40–43). We next examined if either of these transcription factors was Wnt-activated. Indeed, in our pancreatic cancer transcriptomic dataset, both *FOXM1* and *MYBL2* were robustly Wnt-activated. Expression of both *MYBL2* and *FOXM1* was decreased by ∼90% during PORCN blockade (Figures 5B and S2A). Consistent with this, PORCN inhibition decreased MYBL2 protein abundance in HPAF-II pancreatic tumors (Figure 5C). This decrease in *MYBL2* transcripts as well as protein was abrogated by the expression of stabilized β-catenin (Figures 5D i and ii). Thus, both *FOXM1* and *MYBL2,* implicated in the expression of DNA repair pathways, are Wnt-regulated.

**Figure 5:**
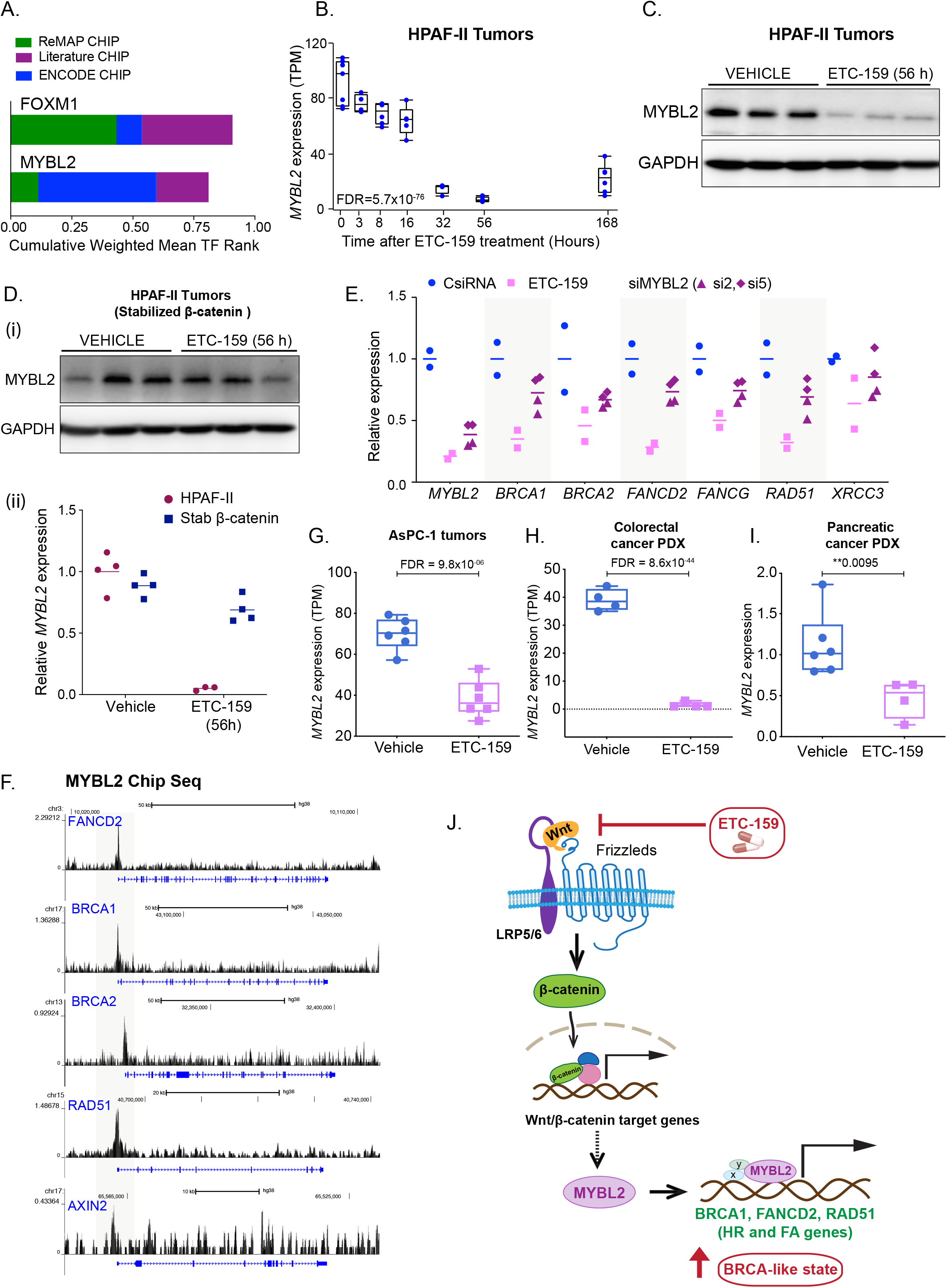
Wnts regulate the expression of HR and FA pathway genes via MYBL2. **A. *Wnt-activated genes show TFBS enrichment for FOXM1 and MYBL2.*** Promoters of *Wnt-activated* HR and FA pathway genes were scanned for transcription factor binding site (TFBS) motifs using ENCODE, ReMAP, and Literature ChIP-seq databases using the web-based tool, CHEA3. B. Temporal regulation of *MYBL2* in HPAF-II orthotopic xenografts treated with ETC-159. Each data point represents an individual tumor. **C. *MYBL2 protein levels are reduced upon ETC-159 treatment in HPAF-II xenografts.*** Tumor lysates from vehicle and ETC-159-treated mice were resolved by SDS-PAGE followed by immunoblotting with the indicated antibodies. Each lane represents an individual tumor **D. *Expression of MYBL2 is regulated by β-catenin***. Expression of *MYBL2* (i) protein levels as measured by immunoblotting and (ii) transcripts as measured by qRT-PCR in HPAF-II tumors with or without stabilized β-catenin treated with vehicle or ETC-159 for 56 hours. Stabilized β-catenin prevents the ETC-159 induced decrease in both *MYBL2* transcripts and protein levels. **E. *Expression of DNA repair genes in Wnt-addicted cells is regulated by MYBL2.*** HPAF-II cells were transfected with two independent siRNAs against *MYBL2* or treated with ETC-159 (100 nM) for 48 hours. Total RNA was isolated and expression of *MYBL2* and DNA repair genes was measured by qRT-PCR. Data are representative of three independent experiments. F. ChIP-seq data from K562 cells from ENCODE shows the binding of MYBL2 on the promoters of DNA repair genes but not on AXIN2 (44). **G-I. *Expression of MYBL2 is downregulated upon ETC-159 treatment in multiple Wnt-addicted tumors.*** Expression of *MYBL2* in AsPC-1 xenograft tumors and colorectal cancer PDX (as measured by RNA-seq) or in pancreatic cancer PDX with *RNF43* mutation (measured with qRT-PCR) is reduced with ETC-159 treatment. Each data point represents an individual tumor. n = 3-5 tumors/group. **J. *PORCN inhibitor ETC-159 creates a BRCA like state:*** Wnt ligand binding to the cell surface receptors Frizzled (Fzd) and LRP5/6 leads to the activation of Wnt/β-catenin target genes such as *MYBL2* that in turn activates the HR and FA pathway genes. ETC-159 treatment inhibits this HR and FA pathway and consequently enhances sensitivity to olaparib.

We assessed the relative importance of *MYBL2* and *FOXM1* in the regulation of HR and FA genes in the Wnt-addicted cells. Knockdown of *FOXM1* with two independent siRNAs had no effect on HR gene expression (Figure S2B). However, knockdown of *MYBL2* in HPAF-II cells in culture using two independent siRNAs reduced the expression of HR and FA pathway genes by 30-50% (Figure 5E). Supporting the role of MYBL2 in regulation of DNA repair genes, we observed that binding sites for MYBL2 were enriched in the promoters of DNA repair genes but not on the promoters of direct Wnt target genes such as *AXIN2* (Figure 5F) (44). Further, examining our previously published datasets (27, 28) we observed that, similar to the regulation of DNA repair genes, *MYBL2* expression was regulated by Wnt signaling in AsPC-1 orthotopic xenografts, a pancreatic cancer PDX with an RNF43 mutation, and in a colorectal PDX with an R-spondin translocation (Figures 5G-I). Taken together, this indicates that the expression of HR and FA genes is in part regulated by a WNT/β-catenin/MYBL2 axis as shown in a model figure 5J.

Finally, we examined if the Wnt target gene *MYC* played a role in the regulation of the HR and FA genes in Wnt high tumors (45–47). We previously examined HPAF-II tumors containing ectopically expressed stabilized MYC (T58A) where PORCN inhibition did not decrease MYC protein abundance (28). Both the basal expression of HR and FA pathway genes and their response to ETC-159 treatment was only modestly influenced by the presence of stabilized MYC (Figure S2C) suggesting that the regulation of HR and FA pathways is not via a Wnt/MYC axis.

### Olaparib and ETC-159 combination enhances senescence

We examined the mechanism by which combination of Wnt and PARP inhibition caused tumor growth inhibition. PARP inhibition in BRCA1 deficient cancers has been reported to cause apoptosis as well as senescence (48, 49). We analyzed the HPAF-II xenografts described in Figure 1C-D, following treatment with the combination of the PORCN and PARP inhibitors. Olaparib alone had no effect on the abundance of HR and FA pathway proteins while ETC-159 alone and in combination with olaparib reduced the protein levels (Figure 6A). Unexpectedly, neither drug alone, nor in combination, caused an increase in markers of apoptotic cell death. There was no increase in cleaved caspase 3 or cleaved PARP in tumors treated with either ETC-159 or olaparib alone, or in tumors treated with a combination of these two inhibitors, showing that synergistic growth inhibition was not working via enhancing cell death by apoptosis (Figure 6B).

**Figure 6:**
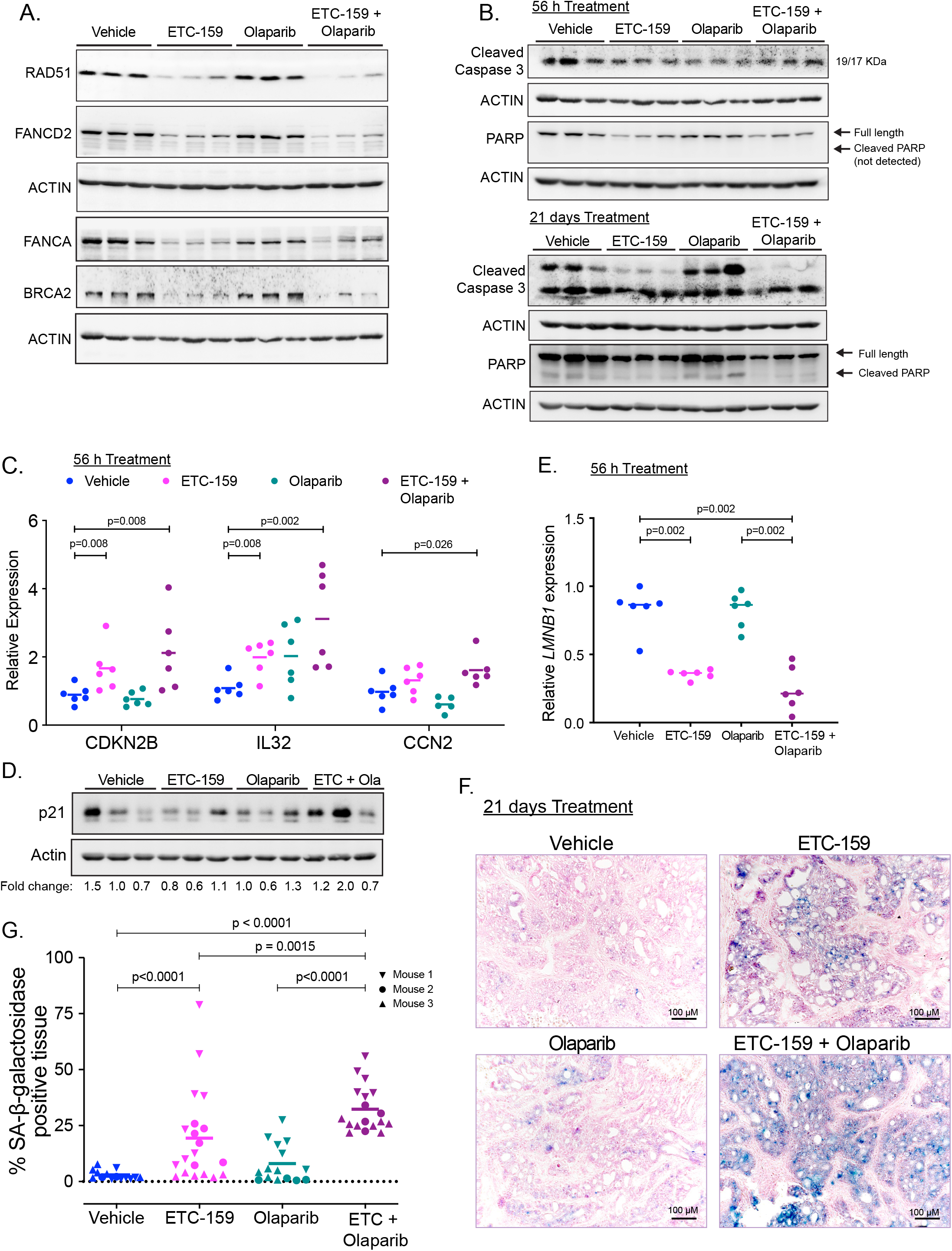
Combination of Wnt inhibition and Olaparib enhances senescence. **A. *Protein levels of HR and FA pathway genes are reduced upon Wnt inhibition in both ETC-159 treated and combination groups.*** Tumor lysates from mice treated with vehicle, ETC-159, olaparib or combination of ETC-159 and olaparib (as shown in Figure 1C) for 56 hours were resolved on a 10% SDS-PAGE gel. The indicated HR and FA pathway proteins were detected by immunoblotting. Each lane represents an individual tumor. **B. *Wnt inhibition alone or in combination with olaparib does not induce apoptosis***. Tumor lysates from mice treated as above for 56 hours or 21 days were resolved on 12% SDS-PAGE gel and analyzed by immunoblotting with the indicated antibodies. Each lane represents an individual tumor. **C-E. *Co-treatment with olaparib and Wnt inhibitor regulates expression of senescence associated genes.*** (C and E) Expression of senescence-associated genes was analyzed in the tumors from all four treatment groups (treated for 56 hours) by qRT-PCR. Each data point represents an individual tumor. p-values for the significant changes as calculated by Mann-Whitney U test are shown. (D) Tumor lysates from indicated groups were immunoblotted for p21. Each lane represents an individual tumor. **F. *Wnt inhibition induces senescence, which is further enhanced by co-treatment with olaparib***. Representative images of tumor sections from the four treatment groups (treated for 21 days) stained for SA-β-galactosidase, a senescence marker, and counterstained with nuclear fast red. Blue color indicates positive staining for senescent cells. **G.** The percentage of SA-β-galactosidase positively stained area (blue) in the representative images from each of the groups is shown. Multiple areas were imaged from three tumor samples per group (indicated as mouse 1, 2 and 3). p-values for the significant changes as calculated by Mann-Whitney U test are shown.

As DNA damaging agents can also induce a senescence response, we next analyzed the expression of genes associated with senescence (50). Among the well-established markers of senescence, *CDKN2A* (encoding 16^INK4a^ and p14^ARF^) is inactive in HPAF-II cells due to a large in-frame deletion (COSMIC database). *CDKN2B* (encoding p15INK^4b^) acts as a tumor suppressor by promoting cell cycle arrest and senescence in the absence of *CDKN2A*, especially in pancreatic cancer (51, 52). The expression of *CDKN2B* was significantly enhanced in tumors from mice treated with ETC-159 alone and further increased in tumors treated with the combination containing olaparib (Figure 6C). In addition, expression of the senescence markers *IL32* and *CCN2* and protein levels of p21 (53–55) were also higher in the tumors from the ETC-159 and olaparib combination group compared to the tumors treated with ETC-159 or olaparib alone (Figures 6C-D). Loss of *LMNB1* (lamin B1) is also associated with senescence (56, 57) and we observed a marked reduction in its expression in the tumors treated with ETC-159 either alone or in combination with olaparib (Figure 6E). Consistent with these changes in the expression of genes associated with senescence, the tumors from the ETC-159 treated group stained positive for senescence-associated β-galactosidase (SA-β-gal), with a significant increase in SA-β-gal staining in the tumors from the ETC-159 and olaparib combination group (Figures 6F-G). Overall our results suggest that combination therapy works in two steps: First, Wnt inhibition alone induces premature senescence. Second, the reduced expression of DNA repair genes following ETC-159 treatment causes HR deficiency that, when combined with blockage of PARP-mediated ssDNA repair by olaparib, leads to an accumulation of dsDNA breaks that accelerates the development of senescence.

### Wnt signaling drives expression of HR and FA pathway genes in the stem cell compartment of the intestines

The above studies show that inhibition of aberrant Wnt signaling markedly decreased the expression of HR and FA pathway genes, leading to a BRCA-like state and acquisition of PARP inhibitor sensitivity in tumors. To determine if physiological Wnt signaling also regulates DNA repair pathways *in vivo*, we examined the expression of the HR and FA pathway genes in intestinal crypts. The small intestine is a highly proliferative tissue that renews every 4-5 days. High Wnt/β-catenin signaling in the intestinal crypts is required for maintenance of the stem cell niche with progressive reduction in Wnt signaling as the cells move to the tips of the villi (58).

In agreement with the presence of a Wnt gradient in the intestine, we observed positive nuclear staining for BRCA1, FANCD2 and RAD51 in cells at the base of the crypts where there is high Wnt signaling. Conversely, Wnt-low villus epithelial cells did not show any staining for these proteins (Figure 7A). Indicating that Wnt signaling promotes the expression of HR and FA pathway genes independent of proliferative state of intestinal cells, the expression of these genes was higher in the slow-cycling stem-cell compartment (crypt base) than in the highly proliferative transit-amplifying compartment. This is also consistent with prior studies showing *Lgr5*+ crypt base columnar cells are radioresistant (1). To ask if increased Wnt signaling is sufficient to increase expression of these HR and FA genes in the crypts we analyzed tissue sections from the intestines of APC^min/+^ mice where the Wnt/β-catenin pathway is activated by loss of APC (adenomatous polyposis coli) function (59). Indeed, we observed robust immunostaining for BRCA1, FANCD2 and RAD51 in the intestinal adenomas (Figure 7B).

**Figure 7:**
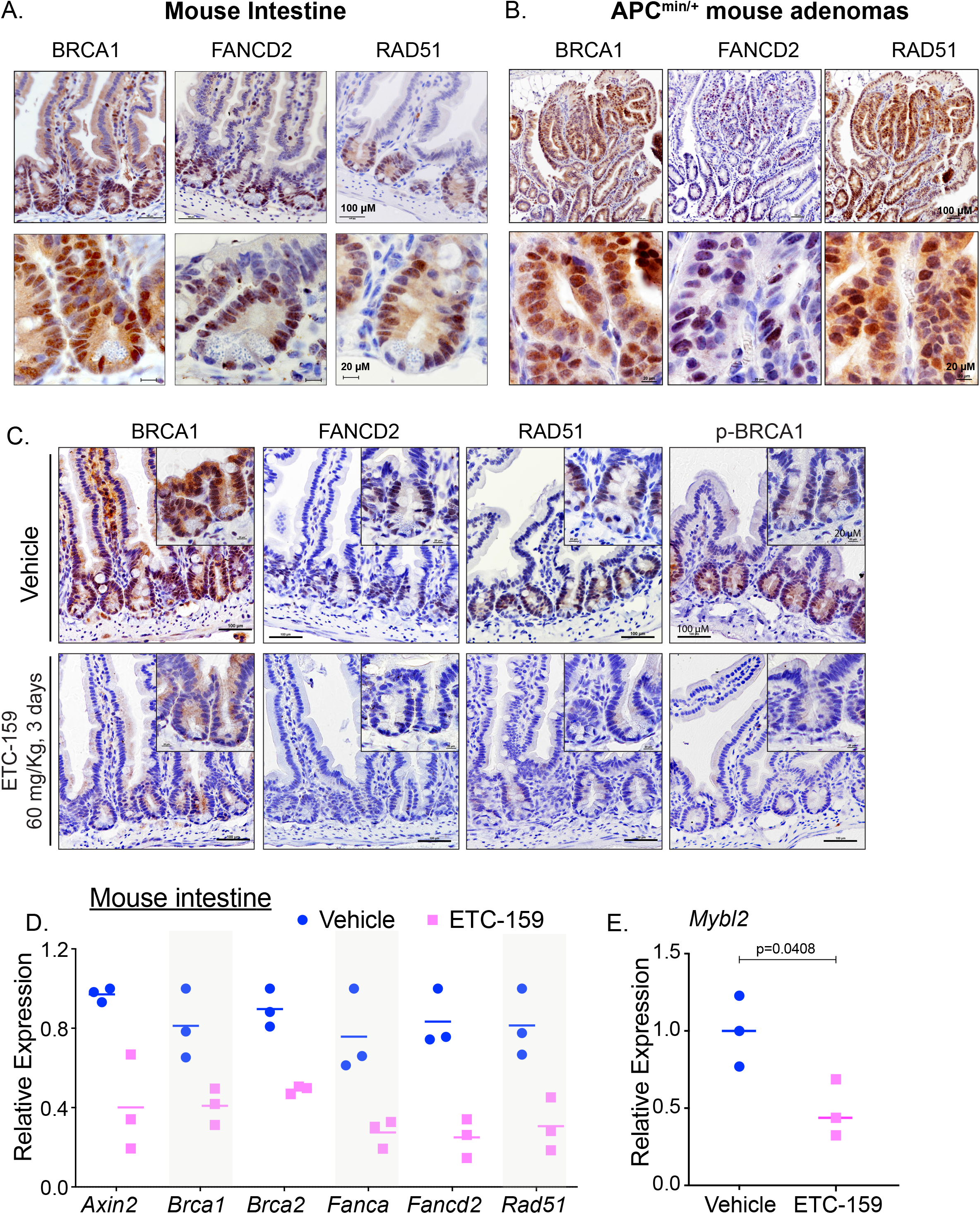
HR and FA pathway genes are expressed in the Wnt high compartment of the small intestine. **A.** Immunohistochemical staining of sections of small intestine from C57BL/6J mice for DNA repair proteins shows that the expression of BRCA1, FANCD2 and RAD51 was high in the crypts (Wnt high compartment) whereas the villi did not show any staining. **B.** Small intestinal adenomas from APC^min/+^ mice displayed positive nuclear immunostaining for HR and FA pathway proteins BRCA1, FANCD2 and RAD51. **C. *PORCN inhibition reduces the expression of HR and FA pathway proteins in the normal mouse intestine.*** C5xs7BL/6J mice were treated with vehicle or ETC-159 (60 mg/kg) for 3 days. Small intestines were sectioned and analyzed by immunostaining for BRCA1, FANCD2, RAD51 and phosphorylated-BRCA1. **D-E. *PORCN inhibition reduces the expression of HR and FA pathway genes and Mybl2 in the normal mouse intestine.*** Total RNA was isolated from small intestines of C57BL/6J mice treated above in Figure 7C. Expression of (D) DNA repair genes and (E) *Mybl2* was measured by qRT-PCR. Each data point represents an individual mouse. n = 3/group

To test if the expression of HR pathway genes in intestinal stem cells is also Wnt-dependent, we treated mice for 3 days with ETC-159. Therapeutically effective doses of ETC-159 are normally well tolerated by mice and do not impact Wnt target genes in the intestine due to the expression of drug pumps by the stromal Wnt-producing cells in the gut. To overcome this intrinsic drug resistance of the gut stroma, we used a high dose (60 mg/kg) of ETC-159 (60). Indeed, Wnt inhibition significantly reduced the expression levels of the HR and FA pathway genes *Brca1*, *Brca2*, *Fanca*, *Fancd2*, and *Rad51* in the intestine. The protein levels of FANCD2, RAD51, BRCA1 and phosphorylated-BRCA1 in the intestinal crypts were also reduced as visualized by immunostaining (Figures 7C-D). Thus, Wnt signaling is necessary for the expression of these genes. Moreover, Wnt inhibition in the normal mouse intestines also lead to reduction in the transcript levels of *Mybl2* in the ETC-159 treated mice (Figure 7E). Taken together this demonstrates that high Wnt signaling regulates DNA damage repair gene expression via Wnt/β-catenin/MYBL2 pathway not only in tumors but also under normal physiological conditions in an adult stem cell compartment.

## DISCUSSION

In this study, we report that Wnt inhibition, via either PORCN or tankyarase inhibition, sensitizes diverse cancers to the PARP inhibitor olaparib. This occurs in part via a Wnt/β-catenin/MYBL2 pathway that enhances the expression of genes involved in homologous recombination and Fanconi anemia pathways, including BRCA1, BRCA2 and multiple FANC genes. We find that Wnt/β-catenin signaling regulates the same genes in the intestinal crypts, suggesting this is a general mechanism by which Wnt/β-catenin signaling can protect the genomes of both Wnt high cancers and adult stem cells.

A genetically defined subset of cancers with high Wnt signaling are vulnerable to Wnt secretion inhibitors. Hence, several small molecules targeting the O-acyltransferase PORCN required for Wnt secretion have advanced to clinical trials. However, cancers invariably acquire resistance to single agent therapies (61). One strategy to overcome drug resistance and reduce adverse effects is synergistic drug combinations. PARP inhibitors like olaparib have successfully moved to the clinics and are now approved treatments of BRCA-mutant ovarian, breast and pancreatic cancers (62–64). The clinical utility of PARP inhibitors is based on the concept of synthetic lethality (7). The enzymes PARP1 and PARP2 are critical for repairing single strand breaks. PARP inhibitors therefore lead to increased unrepaired ssDNA breaks that convert to dsDNA breaks during DNA replication. In the presence of faulty dsDNA break repair, as is seen in BRCA-like states, the accumulated dsDNA breaks cause cell death or senescence (3, 6). Therefore, cancers with loss-of function mutations in HR pathway components such as *BRCA1*, *ATM*, *ATR*, *RAD51* and *FANC genes* are susceptible to treatment with PARP inhibitors (7–9, 32, 65). One key finding of our study was that inhibition of Wnt signaling leads to a coherent downregulation of the expression of *BRCA1/2*, *RAD51*, *FANCD2* and multiple other *FANC* genes, inducing a BRCA-like state in the cells and therefore sensitizing them to the PARP inhibitor olaparib. Tumors develop resistance to PARP inhibitors due to restoration of the HR pathway activity.

Other studies have also suggested Wnt signaling drives enhanced DNA repair. Sun et al. identified WNT16B as a driver of resistance to cytotoxic chemotherapy in prostate cancer (66) while Yamamoto et al. identified WNT3A expression as the cause of in vitro olaparib resistance in an ovarian cancer cell (67). Here we find that Wnts regulate the expression of HR and FA pathway genes in part via a β-catenin/MYBL2 axis. Overexpression of *MYBL2* is observed in several cancers and is associated with poor patient prognosis (68). In p53 mutant cancer cells, *MYBL2* hyperactivation can prevent DNA damage-induced cell cycle arrest (69). MYBL2 is known to regulate cell proliferation, differentiation and tumorigenesis (70, 71). However, its role in regulating the HR and FA pathways has only recently been reported in hematopoietic stem cells (HSCs), where its expression was shown to correlate with the levels of DNA repair genes including RAD51, BRCA1/2 and FANCD2 (42, 43). Low levels of MYBL2 in myelodysplastic syndrome (MDS) patients also preceded transcriptional deregulation of DNA repair genes (72). Our study for the first time shows that Wnt signaling regulates the expression of MYBL2 and this β-catenin/MYBL2 axis regulates expression of HR and FA pathway genes.

Induction of apoptotic cell death is a well-established tumor suppressive response initiated by DNA damaging agents. We did not find any evidence for apoptosis in tumors treated with ETC-159 alone or in combination with olaparib. It has increasingly become clear that several other mechanisms operate to prevent tumor progression. Interestingly, senescence is one of the major outcome following treatment with agents causing DNA crosslinks (48), double-strand breaks (73), as well as PARP inhibition (74). Moreover, depletion of *BRCA1*, *FANCD2* and other *FANC* genes is also linked with premature induction of senescence in response to DNA damaging insults (48, 49, 75, 76). Based on these reports and our results we propose that combination therapy works in two steps. First, ETC-159 treatment inhibits Wnt/β-catenin/MYBL2 pathway leading to HR deficiency, and this, combined with blockage of PARP-mediated ssDNA repair in proliferating cells leads to accumulation of dsDNA breaks. Second, ETC-159 alone induces premature senescence, and this is further enhanced by sustained defective DNA damage response caused by olaparib treatment. Consistent with this, MYBL2 has been shown to inhibit senescence in fibroblasts and cervical cancer HeLa cells (77–79).

A role for Wnt signaling in the control of DNA repair is additionally supported by the high expression of DNA repair genes in the intestinal stem cell compartment where Wnt signaling is high (58). Further, treatment of mice with Wnt secretion inhibitors reduced the expression of both Mybl2 and DNA repair genes in the intestinal crypts. Thus, the role of Wnts in regulating HR and FA pathway genes extends beyond cancers. Wnt signaling in normal stem cells may serve to prevent the mutations arising from normal genotoxic stresses that the intestinal epithelium is exposed to. Notably, ETC-159 is well tolerated at therapeutically effective doses without gut toxicity. This is due to the selective expression of drug-efflux pumps in the Wnt-producing intestinal myofibroblasts (60). Therefore a low dose of ETC-159 used in combination treatment with olaparib results in tumor-selective inhibition of DNA repair pathways while sparing the intestinal stem-cells.

Cells maintain genomic integrity by repairing DNA lesions arising from intracellular processes, such as DNA replication, and extrinsic factors, such as radiation and chemotherapeutic drugs (6). Activation of Wnt signaling is also one of the mechanisms that promotes radioresistance in cancers such as glioblastoma, colorectal, esophageal and nasopharyngeal cancers. This was mainly attributed to the ability of Wnts in promoting cellular proliferation and maintenance of stem cells (12–14, 80). Our study suggests that activation of Wnt signaling in these cancers may enhance their ability to repair DNA damage and hence induce resistance and maintenance of stem cells.

Our findings that inhibition of Wnt signaling regulates DNA repair pathways, specifically the high-fidelity homologous recombination pathway for repairing double-strand breaks, has several implications. It improves our understanding of how high Wnt signaling maintains stemness, provides options for treatment of Wnt high cancers by combination of Wnt inhibitors with PARP inhibitors, and options for overcoming resistance to radiation and chemotherapies by inhibiting Wnt signaling.

## MATERIAL AND METHODS

### Mice, Reagents and Cell Lines

Mice were purchased from InVivos (Singapore) or Jackson Laboratories (Bar Harbor, ME, USA). The Duke-NUS Institutional Animal Care and Use Committee approved all animal studies. Animals were housed in standard cages and were allowed access *ad libitum* to food and water.

ETC-159 was manufactured by Experimental Therapeutics Centre, A*STAR, Singapore. Olaparib (Cat #O-9201) was purchased from LC Laboratories, Woburn MA, 01801. G007-LK purchased from LC laboratories. HPAF-II, CFPAC-1, AsPC-1 and Panc 08.13 and COLO 320HSR were obtained from ATCC, PaTu8988T and EGI-1 from DSMZ. HPAF-II and EGI-1 cell lines were cultured in MEM media with Earle’s salts and non-essential amino acids (Gibco #10370021), AsPC-1 and COLO 320HSR cells in RPMI (Nacalai Tesque #0517625), CFPAC-1 cells in Iscove’s Modified Dulbecco’s Medium (StemCell Technologies #36150) and PaTu8988T cells in DMEM media (Nacalai Tesque #0845935). All media were supplemented with 1 mM sodium pyruvate (Lonza #BW13-115E), 2 mM L-glutamine (Gibco #25030081), 10% FBS (Hyclone SV30160.03) and 1% penicillin/streptomycin antibiotics (Gibco #15140122) and maintained on adherent cell culture flasks in a humidified incubator at 37°C and 5% CO_2_. Panc 08.13 cells were cultured in RPMI media with 15% FBS, 4 μg/ml insulin (Gibco #12585014) and the above-mentioned supplements. All cell lines were routinely tested to be negative for mycoplasma.

For suspension cell culture, optimized number of cells were seeded per well into 24-well ultra-low attachment plates (Corning Costar #3473). Cell clusters and spheroids suspended in the media were collected by centrifugation and rinsed with 1X PBS before RNA isolation.

### Soft agar colony formation assay and drug combination study

HPAF-II, EGI-1, MCAS, CFPAC-1, PaTu8988T or Panc 08.13 cells were plated in 48-well suspension cell culture plates. For each well, optimized number of cells (2000 - 5000 cells per well) mixed with 0.3% agar (Sigma-Aldrich, A1296) or agarose (Sigma-Aldrich, A9045) in 250 μl complete culture media were layered on top of 300 μl of base layer containing 0.6% agar or agarose in complete culture media. After solidification of the agar layers, 250 μl complete culture media containing small molecule inhibitors or DMSO control was added to each well. The cells were allowed to form colonies for 2-3 weeks depending on the colony forming status of each of the cell lines. The colonies were then stained with MTT (3-(4,5-dimethylthiazol-2-yl)-2,5-diphenyltetrazolium bromide) and counted by GelCount (Oxford Optronix, Abingdon, UK). The ED_50_ (the dose that leads to 50% of the maximum colony formation suppressive effect in soft agar) of the inhibitors was determined for each of the cell lines and is listed in Table S1. For determining synergy, cells were treated with either ETC-159, olaparib or a concentration gradient of the combination of the two inhibitors (two replicates per condition). Averages of the total number of colonies obtained as a fraction of the control were then used to determine the Combination Index values using the Chou-Talalay CompuSyn Software (http://www.combosyn.com/)(29).

For low density plating studies, 2000 cells were plated in 48-well cell culture plates coated with poly-L-lysine (Sigma-Aldrich, P4707). COLO 320HSR were treated with combinations of G007-LK, olaparib and DMSO control, and cells were allowed to grow for 1–2 weeks depending on the confluency status. The cells were then stained with 0.5% crystal violet solution (Sigma-Aldrich, V5265) in 25% methanol and counted by GelCount. ED_50_ and combination index values were determined using CompuSyn as above.

### Tumor implantation and treatment of mice

Mouse xenografts were established by subcutaneous injection of HPAF-II cells or HPAF-II cells with stabilized β-catenin in NSG mice. 5 × 10^6^ HPAF-II cells resuspended in 50% Matrigel were used for injection. Mice were treated with ETC-159 or olaparib after establishment of tumors. ETC-159 was formulated in 50% PEG 400 (vol/vol) in water and administered by oral gavage at a dosing volume of 10 μL/g body weight. Olaparib formulated in 50% PEG 400 (vol/vol) in water was administered intraperitoneally at a dosing volume of 10 μL/g body weight. Tumor dimensions were measured with a caliper routinely, and the tumor volumes were calculated as 0.5 × length × width^2^. All mice were sacrificed 8 hours after the last dosing.

### Western Blot

Protein lysates were prepared by homogenizing the tumor tissues in 4% SDS. Protein concentration was determined using Pierce BCA protein assay kit (Thermo Scientific #23225). Equal amount of protein (∼40 μg) was loaded and resolved on a 10% or 12% SDS polyacrylamide gel and then transferred to PVDF membranes. The membranes were probed with antibodies against BRCA1 (Cell Signaling Technology #9010, 1:1000 dilution), BRCA2 (Cell Signaling Technology #10741, 1:1000 dilution), RAD51 (Abcam ab133534, 1:1000 dilution), FANCD2 (Abcam ab108928, 1:1000 dilution), FANCA (Proteintech 11975-1-AP, 1:1000 dilution), MYBL2 (Santa Cruz Biotechnology sc-81192, 1:1000 dilution), cleaved Caspase-3 (Cell Signaling Technology #9661, 1:500 dilution) and PARP (BD Pharmingen 556494, 1:500 dilution). Actin (Abcam ab3280, 1:3000 dilution) or GAPDH (Abcam ab8245, 1:2000 dilution) antibodies were used as protein loading controls. HRP conjugated anti-rabbit IgG (Bio-Rad 1706515) or anti-mouse IgG (Bio-Rad 1721011) secondary antibodies were used at 1:10000 dilution. Blots were visualized on ImageQuant LAS 4000 imager (GE Healthcare Life Sciences).

### Flow cytometric analysis

For detecting endogenous Frizzled levels, PaTu8988T or Panc 08.13 cells were seeded into 6-well plates. After 48 hours cells were stained with 1:100 diluted pan-FZD antibody clone F2.A (81). Anti-human Fc fragment APC-conjugated (Jackson Laboratory, #109-135-098) was used as a secondary antibody. The cells were acquired on BD LSRFortessa and analyzed using FlowJo v10 software.

For cell cycle analysis, HPAF-II cells were seeded in 6-well plates and treated with ETC-159 as indicated. Cells were harvested by trypsinization, washed with PBS and pelleted by centrifugation at 300g for 7 minutes. The cells were fixed in prechilled 70% ethanol, treated with RNase (5 μg/ 50 μl), and stained with propidium iodide (15 μg in 300 μl) at 37°C for 30 minutes in dark. The cells were acquired on BD FACScan and analyzed using FlowJo software.

### Immunohistochemistry

Small intestines from C57BL/6J mice were flushed with PBS, longitudinally cut open, fixed with 10% neutral buffered formalin for 24 hours and embedded as swiss-rolls in paraffin. For immunohistochemistry, 5 μm paraffin sections were deparaffinized, rehydrated and boiled in 10 mM Sodium citrate, pH 6.0 solution for 25 minutes. The sections were then rinsed in TBS and blocked with TBS containing 3% BSA and 0.1% Tween-20 for 1 hour at room temperature followed by incubation with primary antibody diluted in the blocking buffer overnight at 4°C. Next day, sections were rinsed in TBS, incubated with 3% H_2_O_2_ for 15 min at room temperature and then incubated with goat anti-rabbit IgG-HRP (Bio-Rad 170-6515) secondary antibody diluted 1:300 in the blocking buffer for 1 hour at room temperature. Staining was visualized with the Liquid DAB+ Substrate Chromogen System (Dako K3468). Primary antibodies used were RAD51 (Abcam ab133534, 1:400 dilution), FANCD2 (Abcam ab108928, 1:80 dilution), BRCA1 (Proteintech 20649-1-AP, 1:100 dilution) and phosphorylated-BRCA1 (Ser1524) (ThermoFisher Scientific PA5-36627, 1:100 dilution). Images were acquired using Nikon Ni-E microscope with DS-Ri2 camera.

### Senescence associated (SA)-β-galactosidase staining

Fresh frozen OCT-embedded tumor tissues were cut into 8 μm thick sections. The sections were fixed in 2% paraformaldehyde with 0.125% glutaraldehyde solution at room temperature for 5 minutes and rinsed with 1X PBS containing 2 mM MgCl_2_. Thereafter, sections were incubated with X-gal containing staining solution from senescence detection kit (Abcam ab65351) according to manufacturer’s instructions. The sections were counterstained with nuclear fast red (NFR) for 5 minutes. Slides were mounted in mounting media containing gelatin and glycerol. Brightfield images were acquired at 10X magnification using Nikon Ni-E microscope with DS-Ri2 camera. Multiple fields were imaged per tissue section from three tumor samples per treatment group. Quantifications were performed using Fiji ImageJ software. Briefly, color deconvolution was performed to segregate the SA-β-galactosidase (blue stain) positive tissue. Image threshold values were calculated and set to measure the percentage area covered by the entire tissue (NFR stained pink regions) and SA-β-galactosidase positive (blue stained) tissue. Percentage of SA-β-galactosidase positive tissue area was calculated and individual values were plotted with GraphPad Prism. Statistical significance was calculated by Mann-Whitney Unpaired 2-tailed t-test and p-value of < 0.05 was considered a significant difference.

### RNA Analysis

Total RNA from small intestines, cell lines or tumors was extracted using the RNeasy kit (Qiagen #74106). For real time quantitative PCR (qRT-PCR) analysis, RNA was converted to cDNA using iScript cDNA synthesis kit (Bio-Rad #1708891), and SsoFast Eva Green Supermix (Bio-Rad #1725205) was used for qRT-PCR. Gene expression was normalized to *EPN1* and *ACTB* for human cell lines, and *Epn1* and *Pgk1* for mouse tissues. The qRT-PCR primers are listed in Table S2.

### DR-GFP homologous recombination assay

AsPC-1 and Panc 08.13 cells were transfected with pDRGFP plasmid (a gift from Maria Jasin; Addgene #26475)(82). Cells were selected with puromycin (1.2 μg/ml) and single cell clones with stably integrated DR-GFP plasmid were generated. Genomic DNA was isolated from the single cell clones using QIAamp DNA mini kit (Qiagen #51304) and qRT-PCR was performed using DR-GFP-CN1 and DR-GFP-CN2 primers (Table S2) to assess the pDRGFP integration. The expression of DR-GFP inserts was normalized to *ZNF80* and *GPR15* genes (Table S2). A HeLa-DRGFP clone (83) with a single copy of pDRGFP was used as a control to determine the copy number in AsPC-1 and Panc 08.13 DR-GFP single cell clones.

AsPC-1 and Panc 08.13 DR-GFP clones were seeded at optimal density in 24 or 12-well plates and treated with DMSO or 100 nM ETC-159 for 48 hours. The cells were transfected with pCBASceI (a gift from Maria Jasin; Addgene #26477) (84) or pcDNA3.2/V5-DEST (control) plasmid using Lipofectamine2000 (Thermo Fisher Scientific, #11668500) After 6 hours, cells were again treated with DMSO or ETC-159. After 2 days, cells were harvested by trypsinization, acquired on BD LSRFortessa and analyzed using FlowJo v10 software for expression of GFP. The percentage of GFP positive cells was used as a measure of cells undergoing homologous recombination.

### RNA-seq analysis

RNA-seq datasets were analyzed, and clustered as described in Madan *et al.* (28). Briefly, reads originating from mouse (mm10) were removed using Xenome, and the remaining human reads aligned against hg38 (Ensembl version 79) using STAR v2.5.2 (85) and quantified using RSEM v1.2.31 (86). Differential expression analysis was performed using DESeq2 (87). The same pipeline was used to process and analyze RNA-seq reads from a CRC PDX model treated with ETC-159 (27). For the clustering of gene expression changes in HPAF-II orthotopic model, all genes differentially expressed over time (DESeq2, false discovery rate (FDR) < 10%) were clustered using GPClust (88).

### Functional enrichment analysis

For the analysis of the *Wnt-activated* genes, Gene Ontology (GO) enrichments were performed using GOStats (89) using all genes differentially expressed (FDR < 10%) as background. Terms with an FDR < 10% were defined as significantly enriched.

## ACKNOWLEDGEMENTS

We acknowledge the assistance of members of the Virshup lab and members of Experimental Therapeutics Centre. We thank Dr. David Hsu at Duke University for useful discussions. We acknowledge the assistance of the vivarium staff including Hock Lee. We thank Ray Dunn for sections of APC^*min/+*^ mouse intestines. We thank Dr Stephane Angers, Dr Sachdev Sidhu and University of Toronto for the pan-FZD antibody. This research is supported in part by the National Research Foundation Singapore and administered by the Singapore Ministry of Health’s National Medical Research Council NMRC/STAR/0017/2013 (to DMV under the STAR Award Program) and NMRC/OFIRG/0055/2017 (to BM under OF-IRG). BM also acknowledges the support of Duke/Duke-NUS collaborative Research Grant 2017/0040.

## Authors contributions

Babita Madan, Enrico Petretto and David M. Virshup designed the study. Babita Madan, Stanley Lim, Siddhi Patnaik, Sugunavathi Sepramaniam and Amanpreet Kaur performed the animal studies and biochemical analysis. Nathan Harmston designed and performed the bioinformatics analysis. Babita Madan, May Ann Lee, Enrico Petretto and David M. Virshup supervised the study. Babita Madan, Amanpreet Kaur, Enrico Petretto and David M. Virshup wrote the manuscript.

## Declaration of Interests

Babita Madan and David M. Virshup have a financial interest in ETC-159. The authors have no other competing interests

## Notes

### Competing Interest Statement

David Virshup and Babita Madan have financial interest in ETC-159 a drug used in this study

## REFERENCES

1. Hua G et al. Crypt Base Columnar Stem Cells in Small Intestines of Mice Are Radioresistant. Gastroenterology 2012;143(5):1266–1276.

2. Ma J, Setton J, Lee NY, Riaz N, Powell SN. The therapeutic significance of mutational signatures from DNA repair deficiency in cancer. Nat Commun 2018;9(1):646.

3. Dietlein F, Thelen L, Reinhardt HC. Cancer-specific defects in DNA repair pathways as targets for personalized therapeutic approaches. Trends Genet. 2014;30(8):326–339.

4. Deans AJ, West SC. DNA interstrand crosslink repair and cancer. Nat. Rev. Cancer. 2011;11(7):467–480.

5. Ceccaldi R, Sarangi P, D’Andrea AD. The Fanconi anaemia pathway: new players and new functions. Nat Rev Mol Cell Biol 2016;17(6):337–349.

6. Jackson SP, Bartek J. The DNA-damage response in human biology and disease. Nature 2009;461(7267):1071–1078.

7. Ashworth A, Lord CJ. Synthetic lethal therapies for cancer: what’s next after PARP inhibitors? Nat Rev Clin Oncol 2018;15(9):564–576.

8. Farmer H et al. Targeting the DNA repair defect in BRCA mutant cells as a therapeutic strategy. Nature 2005;434(7035):917–921.

9. Bryant HE et al. Specific killing of BRCA2-deficient tumours with inhibitors of poly(ADP-ribose) polymerase. Nature 2005;434(7035):913–917.

10. Clevers H, Loh KM, Nusse R. Stem cell signaling. An integral program for tissue renewal and regeneration: Wnt signaling and stem cell control. Science 2014;346(6205):1248012.

11. Metcalfe C, Kljavin NM, Ybarra R, de Sauvage FJ. Lgr5+ Stem Cells Are Indispensable for Radiation-Induced Intestinal Regeneration. Stem Cell 2014;14(2):149–159.

12. Jun S et al. LIG4 mediates Wnt signalling-induced radioresistance. Nat Commun 2016;7(1):e07301.

13. Emons G et al. Chemoradiotherapy Resistance in Colorectal Cancer Cells is Mediated by Wnt/Œ≤-catenin Signaling. Molecular Cancer Research 2017;:molcanres.0205.2017.

14. Zhao Y et al. Wnt signaling induces radioresistance through upregulating HMGB1 in esophageal squamous cell carcinoma. Cell Death Dis 2018;9(4):9.

15. Luo M et al. FOXO3a knockdown promotes radioresistance in nasopharyngeal carcinoma by inducing epithelial-mesenchymal transition and the Wnt/Œ≤-catenin signaling pathway. Cancer Letters 2019;455:26–35.

16. Karimaian A, Majidinia M, Bannazadeh Baghi H, Yousefi B. The crosstalk between Wnt/β-catenin signaling pathway with DNA damage response and oxidative stress: Implications in cancer therapy. DNA Repair 2017;51:14–19.

17. Nusse R, Clevers H. Wnt/β-Catenin Signaling, Disease, and Emerging Therapeutic Modalities [Internet]. Cell 2017;169(6):985–999.

18. Wu J et al. Whole-exome sequencing of neoplastic cysts of the pancreas reveals recurrent mutations in components of ubiquitin-dependent pathways. Proc. Natl. Acad. Sci. U.S.A. 2011;108(52):21188–21193.

19. Bailey P et al. Genomic analyses identify molecular subtypes of pancreatic cancer. Nature 2016;531(7592):47–52.

20. Ryland GL et al. RNF43is a tumour suppressor gene mutated in mucinous tumours of the ovary. J. Pathol. 2013;229(3):469–476.

21. Wang K et al. Whole-genome sequencing and comprehensive molecular profiling identify new driver mutations in gastric cancer. Nat Genet 2014;46(6):573–582.

22. Giannakis M et al. RNF43 is frequently mutated in colorectal and endometrial cancers. Nat Genet 2014;46(12):1264–1266.

23. Assié G et al. Integrated genomic characterization of adrenocortical carcinoma. Nat Genet 2014;46(6):607–612.

24. Seshagiri S et al. Recurrent R-spondin fusions in colon cancer. Nature 2012;488(7413):660–664.

25. Gurney A et al. Wnt pathway inhibition via the targeting of Frizzled receptors results in decreased growth and tumorigenicity of human tumors. [Internet]. Proc. Natl. Acad. Sci. U.S.A. 2012;109(29):11717–11722.

26. Jiang X et al. Inactivating mutations of RNF43 confer Wnt dependency in pancreatic ductal adenocarcinoma. Proc. Natl. Acad. Sci. U.S.A. 2013;110(31):12649–12654.

27. Madan B et al. Wnt addiction of genetically defined cancers reversed by PORCN inhibition. Oncogene 2016;35(17):2197–2207.

28. Madan B et al. Temporal dynamics of Wnt-dependent transcriptome reveal an oncogenic Wnt/MYC/ribosome axis. J. Clin. Invest. 2018;128(12):5620–5633.

29. Chou T-C. Drug combination studies and their synergy quantification using the Chou-Talalay method. Cancer Res. 2010;70(2):440–446.

30. Zhong Z et al. PORCN inhibition synergizes with PI3K/mTOR inhibition in Wnt-addicted cancers. Oncogene 2019;38(40):6662–6677.

31. Armstrong AC, Clay V. Olaparib in germline-mutated metastatic breast cancer: implications of the OlympiAD trial. Future Oncol 2019;15(20):2327–2335.

32. Lord CJ, Ashworth A. BRCAness revisited. Nat. Rev. Cancer. 2016;16(2):110–120.

33. Ibrahim YH et al. PI3K Inhibition Impairs BRCA1/2 Expression and Sensitizes BRCA-Proficient Triple-Negative Breast Cancer to PARP Inhibition. Cancer Discovery 2012;2(11):1036–1047.

34. Saleh-Gohari N. Conservative homologous recombination preferentially repairs DNA double-strand breaks in the S phase of the cell cycle in human cells. Nucl Acids Res 2004;32(12):3683–3688.

35. Davidson G, Niehrs C. Emerging links between CDK cell cycle regulators and Wnt signaling. Trends in Cell Biology 2010;20(8):453–460.

36. Jasin M, Rothstein R. Repair of Strand Breaks by Homologous Recombination. Cold Spring Harbor Perspectives in Biology 2013;5(11):a012740–a012740.

37. Yu J, Virshup DM. Updating the Wnt pathways. Biosci. Rep. 2014;34(5):593–607.

38. Lau T et al. A Novel Tankyrase Small-Molecule Inhibitor Suppresses APC Mutation-Driven Colorectal Tumor Growth. Cancer Res. 2013;73(10):3132–3144.

39. Keenan AB et al. ChEA3: transcription factor enrichment analysis by orthogonal omics integration. Nucl Acids Res 2019;47(W1):W212–W224.

40. Khongkow P et al. FOXM1 targets NBS1 to regulate DNA damage-induced senescence and epirubicin resistance. Oncogene 2014;33(32):4144–4155.

41. Zona S, Bella L, Burton MJ, Nestal de Moraes G, Lam EW-F. FOXM1: an emerging master regulator of DNA damage response and genotoxic agent resistance. Biochim. Biophys. Acta 2014;1839(11):1316–1322.

42. Bayley R et al. MYBL2 Supports DNA Double Strand Break Repair in Hematopoietic Stem Cells. Cancer Res. 2018;78(20):5767–5779.

43. Bayley R et al. MYBL2 Supports DNA Double Strand Break Repair in Hematopoietic Stem Cells. Cancer Res. 2018;78(20):5767–5779.

44. ENCODE Project Consortium. An integrated encyclopedia of DNA elements in the human genome. Nature 2012;489(7414):57–74.

45. Campaner S, Amati B. Two sides of the Myc-induced DNA damage response: from tumor suppression to tumor maintenance. Cell Div 2012;7(1):6.

46. Guerra L et al. Myc Is Required for Activation of the ATM-Dependent Checkpoints in Response to DNA Damage. PLoS ONE 2010;5(1):e8924.

47. Karlsson A et al. Defective double-strand DNA break repair and chromosomal translocations by MYC overexpression. Proc. Natl. Acad. Sci. U.S.A. 2003;100(17):9974–9979.

48. Santarosa M et al. Premature senescence is a major response to DNA cross-linking agents in BRCA1-defective cells: implication for tailored treatments of BRCA1 mutation carriers. Mol Can Ther 2009;8(4):844–854.

49. Sedic M et al. Haploinsufficiency for BRCA1 leads to cell-type-specific genomic instability and premature senescence. Nat Commun 2015;6(1):7505.

50. Sharpless NE, Sherr CJ. Forging a signature of in vivo senescence. Nat. Rev. Cancer. 2015;15(7):397–408.

51. Tu Q et al. CDKN2B deletion is essential for pancreatic cancer development instead of unmeaningful co-deletion due to juxtaposition to CDKN2A. Oncogene 2017;37(1):128–138.

52. Krimpenfort P et al. p15Ink4b is a critical tumour suppressor in the absence of p16Ink4a. Nature 2007;448(7156):943–946.

53. Jun J-I, Lau LF. CCN2 induces cellular senescence in fibroblasts. J Cell Commun Signal 2017;11(1):15–23.

54. Staal FJ et al. Wnt signaling is required for thymocyte development and activates Tcf-1 mediated transcription. Eur. J. Immunol. 2001;31(1):285–293.

55. Hsu C-H, Altschuler SJ, Wu LF. Patterns of Early p21 Dynamics Determine Proliferation-Senescence Cell Fate after Chemotherapy. Cell 2019;178(2):361–373.e12.

56. Freund A, Laberge R-M, Demaria M, Campisi J. Lamin B1 loss is a senescence-associated biomarker. Molecular Biology of the Cell 2012;23(11):2066–2075.

57. Wang AS, Ong PF, Chojnowski A, Clavel C, Dreesen O. Loss of lamin B1 is a biomarker to quantify cellular senescence in photoaged skin. Sci. Rep. 2017;:1–8.

58. van der Flier LG, Clevers H. Stem Cells, Self-Renewal, and Differentiation in the Intestinal Epithelium. Annu. Rev. Physiol. 2009;71(1):241–260.

59. Su LK et al. Multiple intestinal neoplasia caused by a mutation in the murine homolog of the APC gene. Science 1992;256(5057):668–670.

60. Chee YC et al. Intrinsic Xenobiotic Resistance of the Intestinal Stem Cell Niche. Developmental Cell 2018;46(6):681–695.e5.

61. Zhong Z, Virshup DM. Wnt Signaling and Drug Resistance in Cancer. Molecular Pharmacology 2020;97(2):72–89.

62. Moore K et al. Maintenance Olaparib in Patients with Newly Diagnosed Advanced Ovarian Cancer. N Engl J Med 2018;379(26):2495–2505.

63. Robson M et al. Olaparib for Metastatic Breast Cancer in Patients with a Germline BRCA Mutation. N Engl J Med 2017;377(6):523–533.

64. Golan T, Locker GY, Kindler HL. Maintenance Olaparib for Metastatic Pancreatic Cancer. Reply. N Engl J Med 2019;381(15):1492–1493.

65. McCabe N et al. Deficiency in the repair of DNA damage by homologous recombination and sensitivity to poly(ADP-ribose) polymerase inhibition. Cancer Res. 2006;66(16):8109–8115.

66. Sun Y et al. Treatment-induced damage to the tumor micro-environment promotes prostate cancer therapy resistance through WNT16B. Nat Med 2012;18(9):1359–1368.

67. Yamamoto TM et al. Activation of Wnt signaling promotes olaparib resistant ovarian cancer. Mol. Carcinog. 2019;68:7.

68. Musa J, Aynaud M-M, Mirabeau O, Delattre O, Grünewald TG. MYBL2 (B-Myb): a central regulator of cell proliferation, cell survival and differentiation involved in tumorigenesis. Cell Death Dis 2017;8(6):e2895–e2895.

69. Mannefeld M, Klassen E, Gaubatz S. B-MYB is required for recovery from the DNA damage-induced G2 checkpoint in p53 mutant cells. Cancer Res. 2009;69(9):4073–4080.

70. Tarasov KV et al. B-MYB is essential for normal cell cycle progression and chromosomal stability of embryonic stem cells. PLoS ONE 2008;3(6):e2478.

71. Baker SJ et al. B-myb is an essential regulator of hematopoietic stem cell and myeloid progenitor cell development. Proc. Natl. Acad. Sci. U.S.A. 2014;111(8):3122–3127.

72. Yang R-M et al. MYB regulates the DNA damage response and components of the homology-directed repair pathway in human estrogen receptor-positive breast cancer cells. Oncogene 2019;38(26):5239–5249.

73. Robles SJ, Adami GR. Agents that cause DNA double strand breaks lead to p16 INK4a enrichment and the premature senescence of normal fibroblasts. Oncogene 1998;16(9):1113–1123.

74. Fleury H et al. Exploiting interconnected synthetic lethal interactions between PARP inhibition and cancer cell reversible senescence. Nat Commun 2019;10(1):2556–15.

75. Helbling-Leclerc A, Dessarps-Freichey F, Evrard C, Rosselli F. Fanconi anemia proteins counteract the implementation of the oncogene-induced senescence program. Sci. Rep. 2019;9(1):17024–11.

76. Zhang X, Sejas DP, Qiu Y, Williams DA, Pang Q. Inflammatory ROS promote and cooperate with the Fanconi anemia mutation for hematopoietic senescence. Journal of Cell Science 2007;120(Pt 9):1572–1583.

77. Masselink H, Vastenhouw N, Bernards R. B-myb rescues ras-induced premature senescence, which requires its transactivation domain. Cancer Letters 2001;171(1):87–101.

78. Martinez I, Cazalla D, Almstead LL, Steitz JA, DiMaio D. miR-29 and miR-30 regulate B-Myb expression during cellular senescence. Proc. Natl. Acad. Sci. U.S.A. 2011;108(2):522–527.

79. Mowla SN, Lam EW-F, Jat PS. Cellular senescence and aging: the role of B-MYB. Aging Cell 2014;13(5):773–779.

80. Kim Y et al. Wnt activation is implicated in glioblastoma radioresistance. Lab Invest 2012;92(3):466–473.

81. Steinhart Z et al. Genome-wide CRISPR screens reveal a Wnt-FZD5 signaling circuit as a druggable vulnerability of RNF43-mutant pancreatic tumors. Nat Med 2017;23(1):60–68.

82. Pierce AJ, Johnson RD, Thompson LH, Jasin M. XRCC3 promotes homology-directed repair of DNA damage in mammalian cells. Genes & Development 1999;13(20):2633–2638.

83. Parvin J, Chiba N, Ransburgh D. Identifying the effects of BRCA1 mutations on homologous recombination using cells that express endogenous wild-type BRCA1. JoVE [published online ahead of print: February 17, 2011];(48). doi:10.3791/2468

84. Richardson C, Moynahan ME, Jasin M. Double-strand break repair by interchromosomal recombination: suppression of chromosomal translocations. Genes & Development 1998;12(24):3831–3842.

85. Dobin A et al. STAR: ultrafast universal RNA-seq aligner. Bioinformatics 2013;29(1):15–21.

86. Li B, Dewey CN. RSEM: accurate transcript quantification from RNA-Seq data with or without a reference genome. BMC Bioinformatics 2011;12(1):323.

87. Love MI, Huber W, Anders S. Moderated estimation of fold change and dispersion for RNA-seq data with DESeq2. Genome Biology 2014;15(12):31–21.

88. Hensman J, Lawrence ND, Rattray M. Hierarchical Bayesian modelling of gene expression time series across irregularly sampled replicates and clusters. BMC Bioinformatics 2013;14(1):252.

89. Falcon S, Gentleman R. Using GOstats to test gene lists for GO term association. Bioinformatics 2007;23(2):257–258.

