## Supplementary Information for "WNT inhibition creates a BRCA-like state in Wnt-addicted cancer"

### Figure S1: (accompanying Figure 2)

**A.** ***Time series analysis clusters genes into distinct patterns based on their transcriptional response to PORCN inhibition.*** Reanalysis of data from (28), where HPAF-II cells were orthotopically injected into the tail of the pancreas. Tumors were established over a period of 28 days and mice were treated with ETC-159 (37.5 mg/kg *bid*). RNA was isolated from the tumors at the indicated time-points and analysed by RNA-seq. The heatmap shows all genes that were differentially expressed over time (FDR < 10%) following PORCN inhibition, clustered into 64 clusters based on their pattern of transcriptional response. The clusters of genes that are robustly downregulated following Wnt inhibition are highlighted with colors and indicated as *Wnt-activated* genes.

**B.** ***Wnt inhibition does not alter the cell cycle phases in HPAF-II cells.*** HPAF-II cells were treated with DMSO or ETC-159 ($IC_{50}$= 3 nM) for 48 hours. After treatment cells were stained with propidium iodide and analysed using flow cytometry to determine the number of cells in $G_1$, S or $G_2$/M phase of the cell cycle. Each bar represents mean +/- SD of two replicates.

**C.** ***Wnt inhibition reduces the expression of HR and FA pathway genes in HPAF-II cells.*** HPAF-II cells were treated with DMSO or ETC-159 (100 nM) for 48 hours. Total RNA was isolated, and the normalized expression of DNA repair genes as measured by RNA-seq is shown.

**D.** ***Wnt inhibition reduces the expression of HR and FA pathway genes in Wnt high EGI-1 cells.*** EGI-1 cells were cultured in low adherence plates and treated with DMSO or ETC-159 (100 nM) for 72 hours. Total RNA was isolated, and the expression of *Axin2* and DNA repair genes was measured by qRT-PCR.

Figure S1

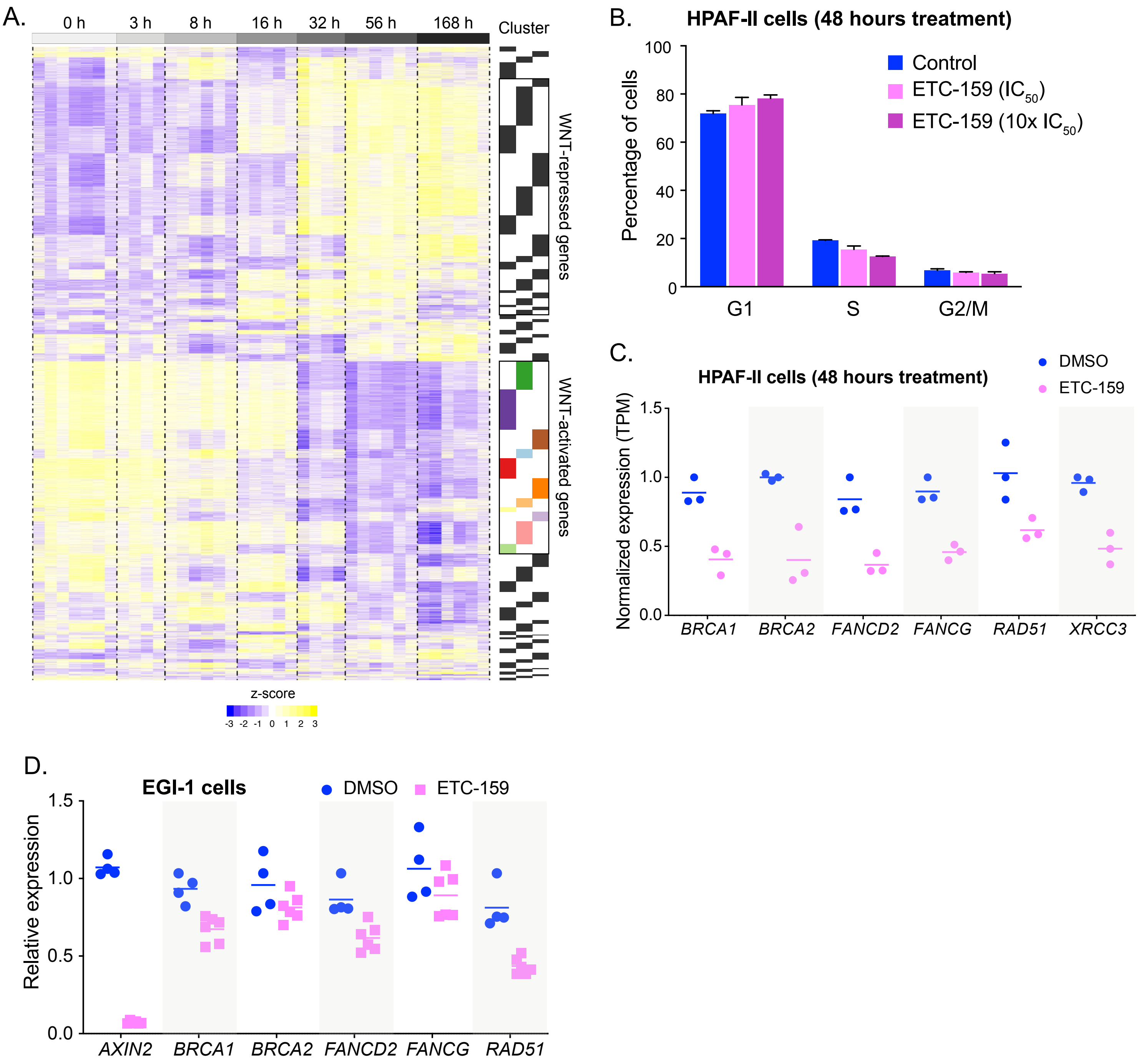
A.
0 h
3 h
8 h
16 h
32 h
56 h
168 h
Cluster
WNT-repressed genes
WNT-activated genes
z-score
-3 -2 -1 0 1 2 3
B.
HPAF-II cells (48 hours treatment)
Control
ETC-159 (IC50)
ETC-159 (10x IC50)
Percentage of cells
100
80
60
40
20
0
G1
S
G2/M
C.
HPAF-II cells (48 hours treatment)
DMSO
ETC-159
Normalized expression (TPM)
1.5
1.0
0.5
0.0
BRCA1
BRCA2
FANCD2
FANCG
RAD51
XRCC3
D.
EGI-1 cells
DMSO
ETC-159
Relative expression
1.5
1.0
0.5
0.0
AXIN2
BRCA1
BRCA2
FANCD2
FANCG
RAD51

### Figure S2: (accompanying Figure 5)

**A.** Temporal regulation of *FOXM1* in HPAF-II orthotopic xenografts treated with ETC-159. Each data point represents an individual tumor.

**B.** ***Expression of DNA repair genes in Wnt addicted cells is minimally regulated by FOXM1.*** HPAF-II cells were transfected with two independent siRNAs against *FOXM1* or treated with ETC-159 (100 nM) for 48 hours. Total RNA was isolated and expression of *FOXM1* and DNA repair genes was measured by qRT-PCR.

**C.** ***Wnt regulated HR and FA pathway gene expression is MYC-independent.*** Mice bearing HPAF-II xenografts without or with stabilized MYC (T58A) were treated with ETC-159 for 56 hours. Tumors from the control and treated groups were harvested and the expression of DNA repair genes was measured.

Figure S2

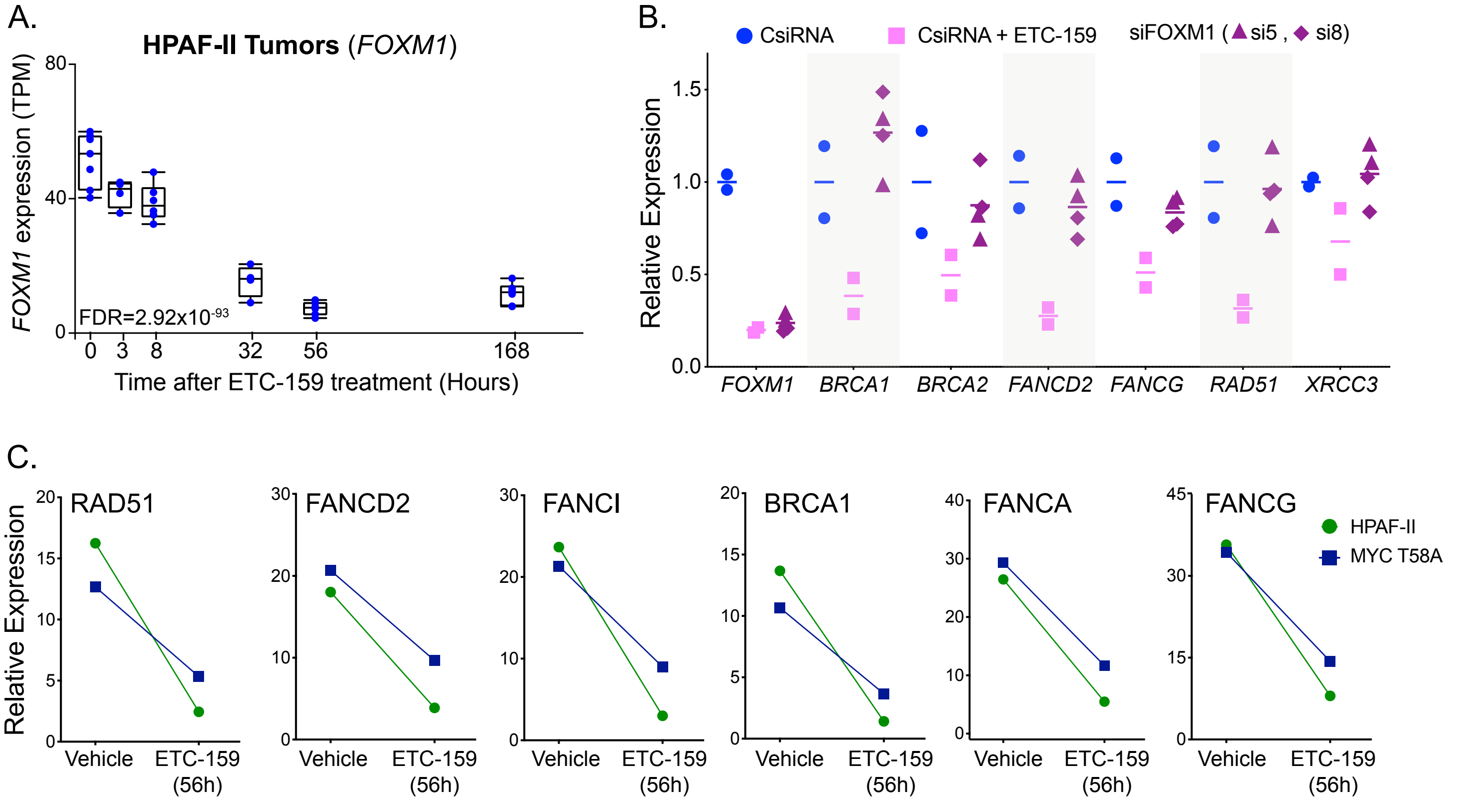
A.
HPAF-II Tumors (FOXM1)
FOXM1 expression (TPM)
80
40
0
FDR=2.92x10-93
0
3
8
32
56
168
Time after ETC-159 treatment (Hours)
B.
CsiRNA
CsiRNA + ETC-159
siFOXM1 (si5 , si8)
Relative Expression
1.5
1.0
0.5
0.0
FOXM1
BRCA1
BRCA2
FANCD2
FANCG
RAD51
XRCC3
C.
Relative Expression
RAD51
FANCD2
FANCI
BRCA1
FANCA
FANCG
HPAF-II
MYC T58A
Vehicle
ETC-159 (56h)

**Table S1:** $IC_{50}$ of the indicated drugs for the respective cell lines in soft agar or low density plating assays.

| Cell Line | **ETC-159 $ED_{50}$** | **Olaparib $ED_{50}$** |
|---|---|---|
| HPAF-II | 0.0077 µM | 39.19 µM |
| EGI-1 | 0.0275 µM | 8.65 µM |
| MCAS | 0.287 µM | 3.979 µM |
| CFPAC-1 | 0.027 µM | 6.25 µM |
| PaTu8988T | 1.1 µM | 1.01 µM |
| COLO 320HSR | 0.5 µM (G007-LK) | 2.0 µM |

**Table S2**: List of qRT-PCR primers

| Gene | Species | Forward primer (5'-3') | Reverse primer (5'-3') |
|---|---|---|---|
| Axin2 | Human, Mouse | CTCCCCACCTTGAATGAAGA | TGGCTGGTGCAAAGACATAG |
| BRCA1 | Human | CAACATGCCCACAGATCAAC | ATGGAAGCCATTGTCCTCTG |
| BRCA2 | Human | CAGAAGCCCTTTGAGAGTGG | TCCATCTGGGCTCCATTTAG |
| FANCD2 | Human | CCCTGAGCTGCTTTTCTTGC | CGGCTTCCTTTGTTCTTGAG |
| FANCG | Human | CTGTTCTTCCCTTGGAGCTG | TCTCTAGGCTCCGCTGGATA |
| RAD51 | Human | TTTGGAGAATTCCGAACTGG | TACATGGCCTTTCCTTCAC |
| XRCC3 | Human | GTGCATCAACCAGGTGACAG | TTAGCCCAGGTTATGCCAAG |
| CTNNB1 | Human | ATGGCTTGGAATGAGACTGCT | CCCATCAACTGGATAGTCAGC |
| MYBL2 | Human | GAATTCCCGAAGCGTGAGGA | CAGGGTCCGACTCGATCAAG |
| FOXM1 | Human | GAGCAGCGACAGGTTAAGGT | GTCATGCGCTTCCTCTCAGT |
| CDKN2B | Human | GGAAAGAAGGGAAGAGTGTCGTT | CGCGCATTCCGCAGC |
| LMNB1 | Human | GATTGCCCAGTTGGAAGCCT | TGGTCTCGTTAATCTCCTCTTCATACA |
| IL32 | Human | TGGCGGCTTATTATGAGGAGC | CTCGGCACCGTAATCCATCTC |
| CCN2 | Human | TTGGCCCAGACCCAACTATG | CAGGAGGCGTTGTCATTGGT |
| ACTB | Human | ATAGCACAGCCTGGATAGCAACGTAC | CACCTTCTACAATGAGCTGCGTGTG |
| EPN1 | Human | CTCTGACTTTGACCGACTCC | TGACCCCACTCATGTCAAAC |
| Brca1 | Mouse | CCGGATACGAGAGTGAAACAA | TGCTGCAGCTTTATCAGGTT |
| Brca2 | Mouse | AGGAAATGTTGGCTGTGTGGA | CGCTGTGTTGTGTCTTCTTCG |
| Fancd2 | Mouse | CAAAATCAGCTAGGTGTGGATCA | CCAGGCCATTAACAAACTCTTCT |
| Fanca | Mouse | GTGGTCGGTGGATGAGATGTT | CCTAACTCCTCTCCACGCAAA |
| Rad51 | Mouse | AAGTTTTGGTCCACAGCCTATTT | CGGTGCATAAGCAACAGCC |
| Mybl2 | Mouse | GTGAGGCAGTTTGGACAGCAA | GGATTCAAAACCCTCAGCCA |
| Pgk1 | Mouse | TCAAAAGCGCACGTCTGCCG | AAGTCCACCCTCATCACGACCC |
| Epn1 | Mouse | TTGTGAGTCGCGTCATTTCTC | CCTCTGAGTAGTTGTGGACGATA |
| DRGFP-CN1 | - | AGATCCGCCACAACATCGAG | TCTCGTTGGGGTCTTTGCTC |
| DRGFP-CN2 | - | AGCAAAGACCCCAACGAGAA | TCGTCCATGCCGAGAGTGAT |
| ZNF80 | Human | CTGTGACCTGCAGCTCATCCT | TAAGTTCTCTGACGTTGACTGATGTG |
| GPR15 | Human | GGTCCCTGGTGGCCTTAATT | TTGCTGGTAATGGGCACACA |
